# Spatial confinement shapes organelle architecture and remodeling in axons

**DOI:** 10.64898/2026.07.07.737043

**Authors:** Mihir Relan, Venkata Mallampalli, Benjamin A. Barad, Andrea K.H. Stavoe, M. Neal Waxham

**Affiliations:** Department of Neurobiology and Anatomy, University of Texas Health Science Center at Houston, Houston, TX 77030; Department of Chemical Physiology and Biochemistry, School of Medicine, Oregon Health & Science University, Portland, OR, USA

**Keywords:** Cryo-electron tomography, axonal cytoskeleton, dorsal root ganglion, neural networks, organelle remodeling, spatial confinement

## Abstract

Axons are extended cellular compartments that mediate neuronal connectivity over extraordinary distances, placing unique demands on local organelle organization and trafficking. How these geometric constraints influence organelle architecture remains poorly understood. Here, we used cryo-electron tomography and quantitative morphometric analyses to define the three-dimensional ultrastructural organization of dorsal root ganglion axons in a near-native state. We identify distinct vesicle populations with tightly versus broadly constrained size distributions and reveal a continuum of endolysosomal and autophagic intermediates that highlight the dynamic nature of membrane remodeling in growing axons. Unexpectedly, we observe vesicular structures enclosed within the lumen of the endoplasmic reticulum, suggesting a previously undescribed mechanism of ER membrane remodeling. Across multiple organelle classes, morphology and size are constrained by axonal geometry. This principle is most evident in mitochondria, which undergo dramatic narrowing and remodeling at varicosity – axon boundaries to traverse confined axonal segments. Together, these findings reveal spatial confinement as a fundamental organizing principle of axonal cell biology.

**SUMMARY:** Cryo-electron tomography establishes a quantitative framework for organelle organization in axons. Diverse membrane trafficking pathways, including endolysosomal intermediates and mitochondria, exhibit structural adaptations to axonal geometry, identifying spatial confinement as a fundamental organizing principle of axonal cell biology.

## INTRODUCTION

Neurons are post-mitotic and exhibit a complex architecture that varies widely across the nervous system. In most neurons, dendrites and axons extend from the cell body and are generally described as receiving and distributing information, respectively. The length and complexity of neuronal processes place unique demands on the organellar systems responsible for maintaining neuronal health (Attwell and Laughlin, 2001; Harris et al., 2012). Axons can exceed a meter in length while often maintaining diameters of only a few hundred nanometers, creating an extreme aspect ratio that poses substantial challenges for intracellular transport, metabolism, and organelle maintenance (Sheng, 2017). Axonal homeostasis is particularly dependent on long-range transport and organelle quality control. Organelles, vesicles, and cytoskeletal components synthesized in the soma must be transported to distal sites via motor proteins along polarized microtubule tracks. Failure of this transport processes contribute to a broad range of peripheral neuropathies and neurodegenerative diseases (Misgeld and Schwarz, 2017).

Along axons, focal swellings termed varicosities represent specialized structural compartments that accumulate organelles. Varicosities are also sites of neurotransmitter and neuropeptide release via en passant secretion along the mature presynaptic compartment. They host mitochondria, vesicular pools, endosomal machinery, endoplasmic reticulum (ER), and cytoskeletal elements (Fischer et al., 2018; Lapios et al., 2025). Recent studies suggest that these variations in axonal geometry are not merely structural features. Watanabe and colleagues demonstrated that nanoscale varicosities emerge from membrane mechanical properties and influence action potential conduction velocity by altering local axonal dimensions (Griswold et al., 2024). Together, these findings indicate that axonal geometry influences both electrical signaling and the intracellular environment through which organelles must be transported. Defining the organization of organelles within varicosities is therefore essential for understanding how the physical constraints imposed by axonal architecture influence neuronal function and homeostasis.

Dorsal root ganglia (DRG) comprise the sensory ganglia of the spinal cord, and their neurons extend axons throughout the body to transmit somatosensory information to the central nervous system. DRG neurons possess several features that make them particularly well suited for investigating axonal organelle organization and trafficking. Dissociated DRG neurons can be maintained in culture and have become a widely-used model for examining axonal transport, autophagy, and protein quality control using high-resolution live-cell imaging (Stavoe et al., 2019; Maday and Holzbaur, 2014). Unlike neurons cultured from the mammalian brain, DRG neurons extend only axonal processes *in vitro*, eliminating ambiguity between dendritic and axonal compartments. In addition, DRG neurons can be isolated from animals across the lifespan, enabling investigation of age-dependent changes in axonal biology (Stavoe et al., 2019). Together, these properties make DRG neurons a powerful model for examining how organelle organization and trafficking adapt to the unique physical constraints of the axonal compartment.

Despite extensive studies of axonal transport and organelle dynamics, a quantitative three-dimensional framework linking organelle architecture, spatial organization, and the physical constraints imposed by axonal geometry remains lacking. DRG neurons are particularly well suited for addressing this gap because they can be cultured directly on EM-compatible substrates and cryopreserved for nanoscale structural analysis by cryo-electron tomography (cryo-ET) (Foster et al., 2021; Shahmoradian et al., 2014; Waxham et al., 2026). Cryo-ET permits reconstruction of cellular architecture as three-dimensional volumes in a near-native state without fixation, staining, or other processing steps that can perturb ultrastructure (Turk and Baumeister, 2020; Lučić et al., 2007). Fortuitously, DRG axons and their associated varicosities (0.2–2 µm in diameter) are sufficiently thin to be imaged directly without cryo-sectioning or focused ion beam milling, substantially increasing data acquisition rates (Foster et al., 2021; Waxham et al., 2026). Recent advances in deep-learning-based semantic segmentation and computational morphometrics further enable quantitative measurements of organelle size, morphology, abundance, and spatial relationships within cellular volumes (Chen et al., 2017; Heebner et al., 2022; Lamm et al., 2025; Medina et al., 2026; Barad et al., 2023).

In the present study, we produced a large cohort of cryo-preserved DRG axon tomograms and applied quantitative morphometric analyses to define the organization of membrane-bound organelles within their native cellular context. These analyses establish a quantitative ultrastructural framework for the axonal compartment and reveal how organelles adapt to the physical constraints imposed by axonal geometry. Together, these observations suggest that spatial confinement is a fundamental organizing principle of axonal cell biology, shaping organelle architecture and driving membrane remodeling across multiple trafficking pathways.

## RESULTS

### Axons and varicosities create a heterogeneous landscape of spatial confinement

To define how organelles are organized within the spatially constrained environment of axons, we generated a large cohort of cryo-electron tomograms from primary DRG neurons and quantified the morphology and spatial relationships of membrane-bound organelles. This dataset provides a three-dimensional framework for examining how organelles are distributed within axons and varicosities and establishes the basis for assessing how axonal geometry influences organelle architecture. DRGs from young adult mice (3-month-old) were cultured on gold Quantifoil grids for 48 hr before cryopreservation, as previously described (Waxham et al., 2026). We first collected montage overviews of DRG axons traversing the carbon surface of the cryo-preserved specimens. Within this data, we could easily distinguish varicosities and the axons that interconnect them (Figure 1A). The varicosities are axonal swellings that harbor collections of organelles and cytoskeletal elements, and they form attached to the carbon (Fig. 1Aa) as well as over the holes in the Quantifoil carbon surface (Fig. 1Ab). To provide maximum contrast for high-resolution tomography, we primarily targeted data collection on cryo-preserved varicosities overlying holes in the Quantifoil grids; however, a subset of tomograms from structures on the carbon surface was also collected to ensure observations were not biased by mechanical force on varicosities in holes. From a sampling of 10 randomly selected montages of low-magnification TEM images, the varicosity diameter ranged from 0.6 to 1.88 μm, with a median of 1.27 ± 0.34 μm (Fig. 1B; n = 82). We observed that the diameter of the axons can be quite variable: from 0.07 to 0.38 μm with a median of 0.15 ± 0.06 μm (Fig. 1B, n = 552). Interestingly, the axonal diameter was variable even along a stretch of a single axon segment, indicating that organelles encounter a highly heterogeneous physical environment during transport.

**Figure 1.**
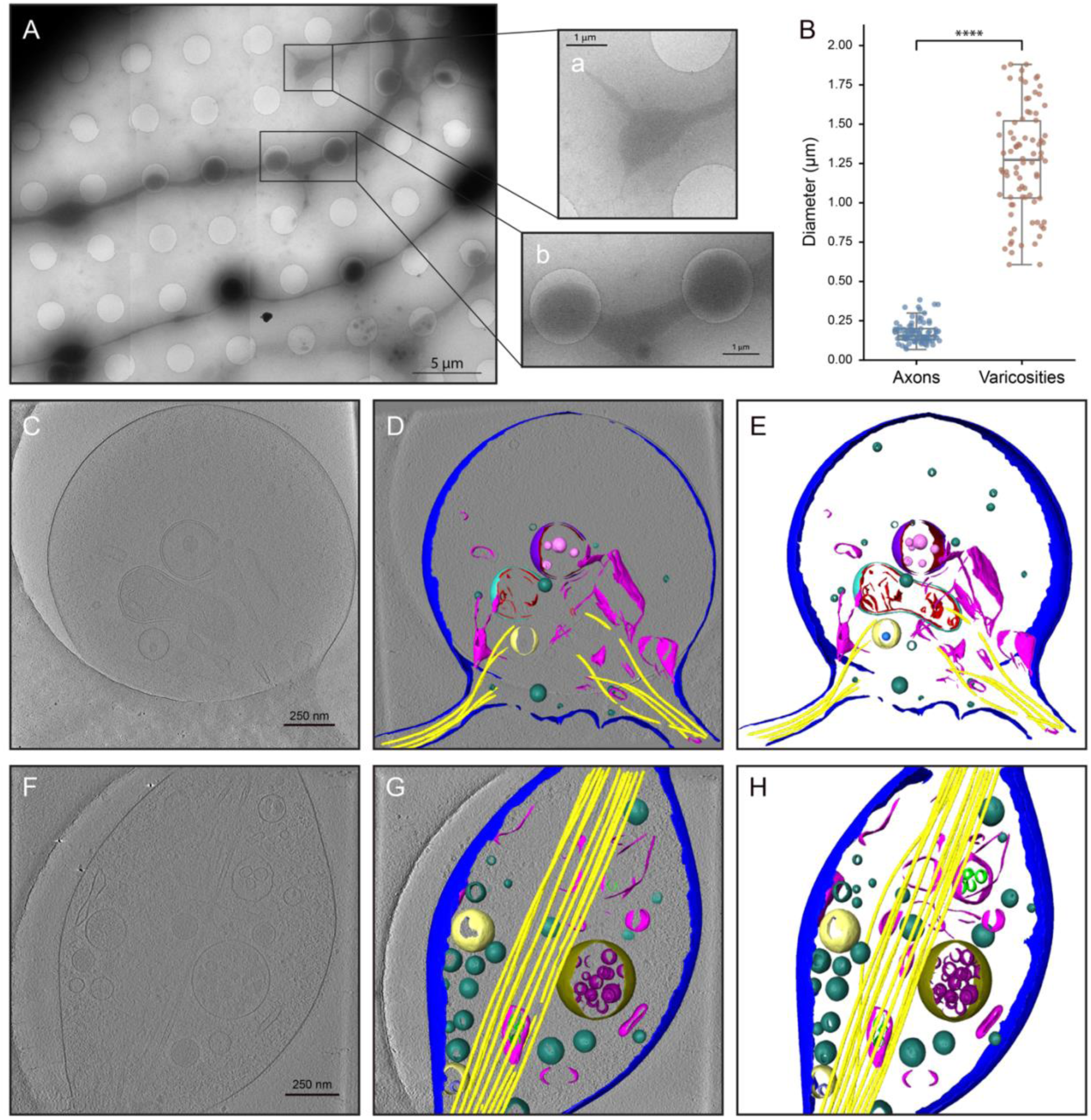
Overview of DRGs cultured on EM grids. **(A)** Cryo-electron micrograph of a low magnification montage showing multiple DRG axonal processes extending across the grid. Visible is the SiO_2_ surface that has 2 μm holes spaced 2 μm apart. Scale bar is 5 μm. Two insets from the indicated boxes in (A) show a process tip growing on the carbon surface (a) and a pair of varicosities suspended in the 2 μm holes of the Quantifoil surface (b). Scale bars on the insets are 1 μm. **(B)** Box plot showing the diameter (in μm) of axons (n= 552 measurements; random data points visualized) and maximum diameter of varicosities (n = 82) from a random collection of 10 micrographs like that shown in (A). Panels **(C and F)** are slices through tomographic reconstructions of two representative tomograms. Scale bars are 250 nm. **(D and G)** are semantic segmentations of the same two tomograms overlaid on the tomographic slice and **(E and H)** are the complete 3D segmentations. In **(D and E)** the plasma membrane (PM) is in blue, endoplasmic reticulum (ER) is in magenta, free vesicles are in light green, microtubules (MTs) are in yellow, phagophore outer membrane is in purple, phagophore inner membrane is in red vesicles internal to the phagophore are in light pink, outer mitochondrial membrane is in light blue, the inner mitochondrial membrane is in red and an endosome is in light yellow. In **(G and H)** the color code is the same as in (D and E) with the addition of the outer membrane of the multi-vesicular body (MVB) in brown, MVB vesicles in dark purple, ER vesicles in green and endosomal vesicles in dark blue.

To characterize the number and types of organelles present, we employed a combination of visual inspection, machine-learning-based segmentation, and morphometric analyses (Figure 1 and Supplementary Movies 1 and 2). We initially distinguished eight classes of structural elements from this dataset: plasma membrane (PM), microtubules (MT), mitochondria, endoplasmic reticulum (ER), multivesicular bodies (MVB), multilamellar vesicles (MLV), phagophores, endosomes, and free vesicles. Some of these vesicular structures contain internal vesicles (Fig. 1E and 1H) that were segmented separately for subsequent analysis: phagophore , MVB , MLV , endosome , and ER . We acknowledge that organelle classes can span a developmental/maturational spectrum and, with some exceptions, it is difficult to assign them solely on morphology (Klumperman and Raposo, 2014; Bieber et al., 2022). Therefore, we applied conservative morphological criteria to assign each organelle to a class and applied them consistently across datasets.

As a global assessment of the organelles present in the dataset, all 192 tomograms were inspected for the presence or absence of eight major organelle classes (Table 1). Free vesicles were abundant, and examples could be identified in every tomogram. ER was also nearly ubiquitous, with one exception (191/192 tomograms). MTs were found in 96% of the tomograms, and endosomes, mitochondria, and MVBs were the next most represented classes (52-53%). The frequency of MLVs was lower (33%), and finally, phagophores were the least frequent class (5%). Detailed characteristics of each identified class are investigated below.

**Table 1.**
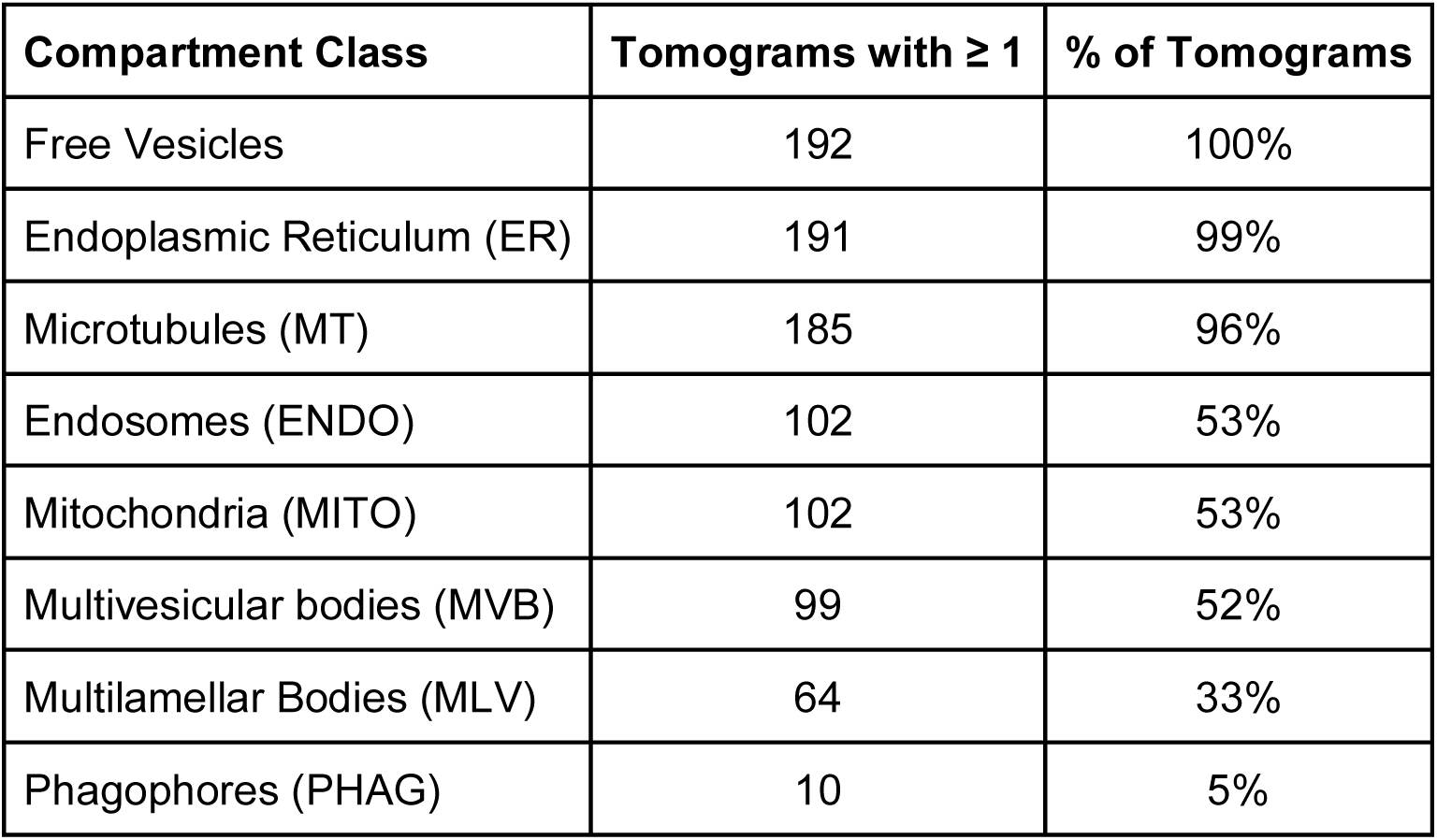
Quantification of organelles in 192 tomograms.

| Compartment Class | Tomograms with $\geq 1$ | % of Tomograms |
| --- | --- | --- |
| Free Vesicles | 192 | 100% |
| Endoplasmic Reticulum (ER) | 191 | 99% |
| Microtubules (MT) | 185 | 96% |
| Endosomes (ENDO) | 102 | 53% |
| Mitochondria (MITO) | 102 | 53% |
| Multivesicular bodies (MVB) | 99 | 52% |
| Multilamellar Bodies (MLV) | 64 | 33% |
| Phagophores (PHAG) | 10 | 5% |

### The cytoskeleton is dominated by microtubules

As expected, MTs were ubiquitous in the varicosities. A previous study undertook an extensive description of MTs and other cytoskeletal elements in cryo-preserved DRG axons prepared similarly to those in the present study (Foster et al., 2021). In our tomograms, MTs were observed tracking linearly through varicosities (Fig. 1H) or occasionally taking more circuitous pathways (Fig. 1E). MTs were nearly ubiquitous across the tomograms but were variable in number. The median number of MTs was 4 per varicosity (Fig. 2A). The average MT diameter was 24.8 ± 1.3 nm, similar to that described previously (21.2 nm) (Foster et al., 2021). The curved nature of the MTs in the varicosities is notable (Fig. 1D, 2C-2E). We calculated the persistence length for the entire population of MTs and found a median of 151 ± 130 µm (Fig. 2F). Additionally, we observed a range of MT bending in varicosities across 1098 MTs (Fig. 2C-2E). Bending was not a feature of all MTs (compare MTs in Fig. 1E to those in Fig. 1H). In vitro measurements place MT persistence length in the mm range (Gittes et al., 1993; Pampaloni et al., 2006; Brangwynne et al., 2007), significantly longer than the curvature observed in our tomograms in DRG varicosities. The apparent persistence length inferred from our measurements is consistent with MTs experiencing mechanically imposed curvature arising from confinement, crosslinking, lattice defects, or other local forces. The MTs form straighter linear arrays when entering and exiting a varicosity, respecting the narrowed constraints imposed by the axonal boundaries (Fig. 1E and 2C).

**Figure 2.**
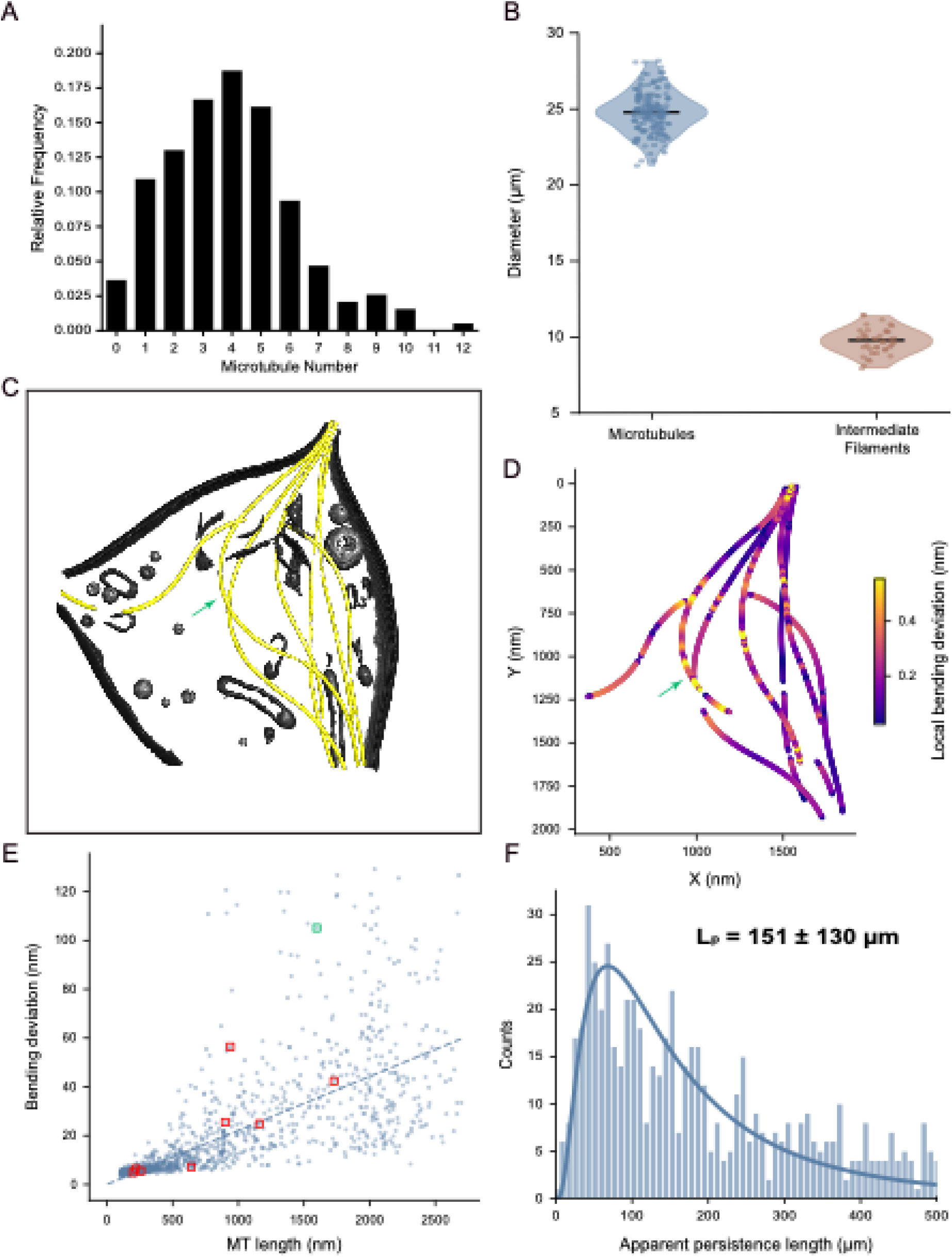
Characteristics of cytoskeletal elements in DRG varicosities. **(A)** The relative frequency of MTs found in DRG varicosities across 192 tomograms demonstrates a median number of MTs as 4 with a range from 0-12. **(B)** A violin plot representing the diameter of MTs and intermediate filaments is shown. The average diameters were 24.8 ± 1.3 nm (n = 171) and 9.70 ± 10.85 nm (n = 40), respectively, indicated by the black horizontal bars. **(C)** Pseudo-colored segmentation of a DRG varicosity with the membranes in various greyscale tones and the MTs highlighted in yellow. **(D)** The segmentation in (C) was subjected to an analysis of local bending deviations (see Methods) that are plotted in pseudo color reflecting the magnitude of the bending deviation as shown on the heatmap on the right. **(E)** Scatter plot of MT instance bending deviation versus MT length (n = 1098). The dotted line passes through the origin with a slope equal to the median normalized bending deviation (0.022). The interpretation is the median MT deviates laterally by 22 nm per 1,000 nm of length. A perfectly straight MT over its length (no bending deviation) would produce a value of zero. The point outlined by green highlights the curved MT identified by a green arrow in (D). The 8 red squares highlight the points representing the other MTs shown in (D). **(F)** Histogram of apparent persistence length across the MT population (n = 598) with lognormal fit. Instances were restricted to MTs with lengths between 200 and 2,000 nm and minimum bending deviation of 1 nm. Median apparent persistence length was 151 ± 130 µm.

Foster et al. identified abundant actin and intermediate filaments (they described as thick filaments) in their tomograms of cryopreserved DRG axons (Foster et al., 2021). While we observed actin and intermediate filaments in a subset of tomograms (Fig. S1), we could not detect actin in most varicosities. One exception was the abundant actin that we could identify in what appeared to be the tips of axons. At the resolution of our tomograms, we could not reliably measure the thickness of the actin filaments which may have contributed to our inability to visualize these thin filaments. However, we could identify a sufficient number of intermediate filaments to enable quantification (Fig. 2B and S1). The average diameter of intermediate filaments was 9.70 ± 0.85 nm (n=40), similar to the diameter of thick filaments (8.5 nm) reported previously (Foster et al., 2021). While insufficient in number to quantify curvature reliably, we observed intermediate filaments exhibiting straight (Fig. S1A) and bent (Fig. S1C, D, and E) morphologies.

### Free vesicles resolve into light and dark lumen populations

Single-membrane “free” vesicles are abundant axonal cargos and have been described in both conventional electron microscopy and cryo-ET studies of neuronal processes (Nakata et al., 1998; Sorra et al., 2006; Zhai et al., 2001; Schrod et al., 2018; Tao et al., 2018). Free vesicles were the most abundant membrane-enclosed structure and were present in every tomographic reconstruction (Fig. 3A and Table 1). We classified free vesicles based on two criteria: sphericity and lack of visible membrane material in their lumen. Two distinct, identifiable vesicle types were obvious: small, light-lumen vesicles and large, dark-lumen vesicles (Fig. 3A). These two vesicle types have been described in numerous previous analyses, including in DRG axons (Foster et al., 2021) and likely include transport vesicles and potential neurotransmitter and neuropeptide-containing vesicles.

**Figure 3.**
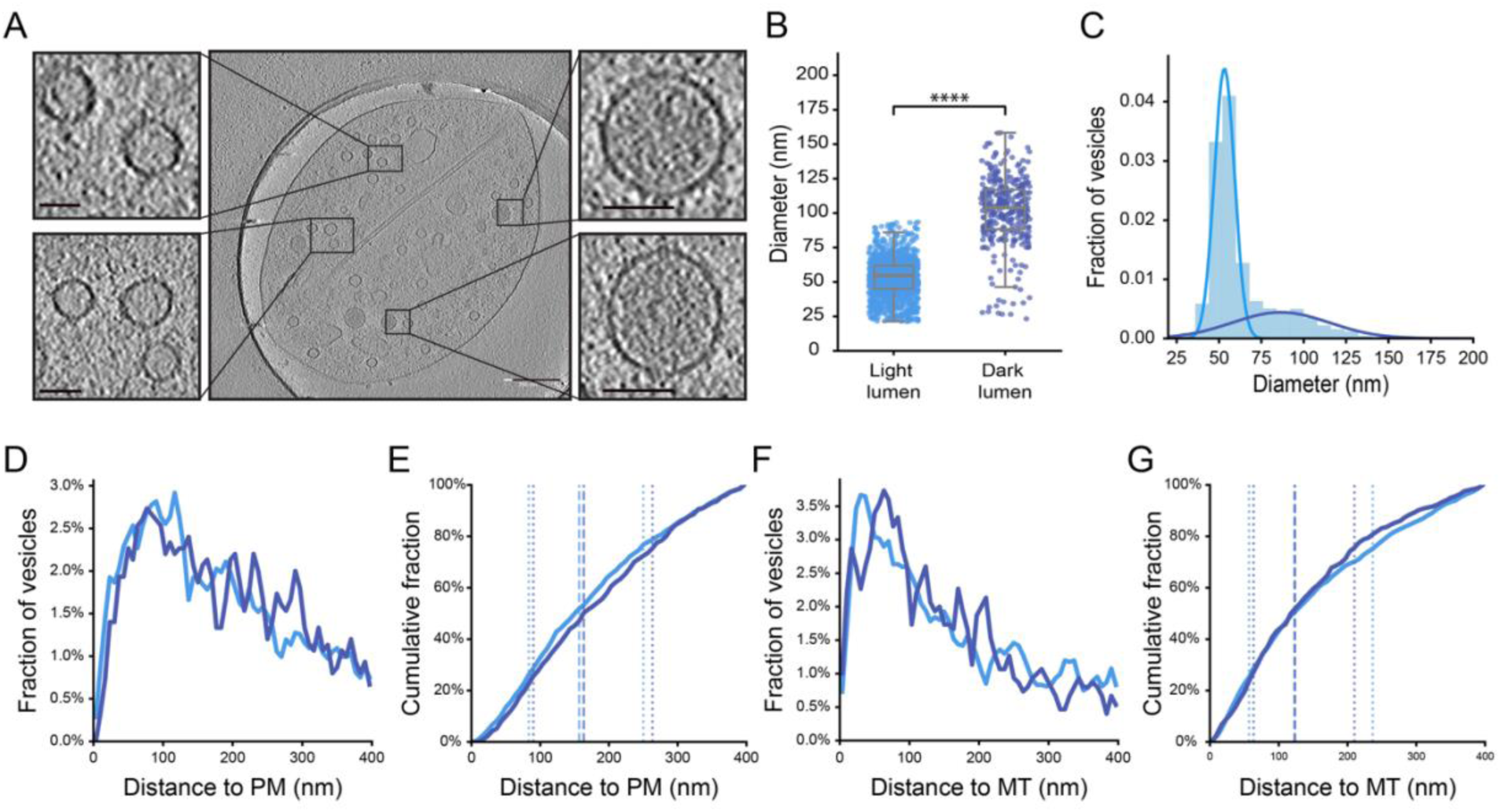
Size and spatial characteristics of free vesicles in DRG varicosities. **(A)** Tomogram slices of spherical light- and dark-lumen vesicles. The blow ups represent a selection of light-lumen vesicles (left) and dark-lumen vesicles (right) from the overview. Scale bar on the overview tomographic slice is 250 nm and scale bars on the insets are 50 nm. **(B)** Diameters of light- (53.8 ± 14.6 nm, n = 1,359) and dark-lumen (102.5 ± 26.5 nm, n = 331) vesicles measured by supervised segmentation. ****, P < 0.0001, Mann-Whitney U test. **(C)** Size distribution of equivalent-sphere diameters for the free vesicle population derived from the Surface Morphometrics pipeline, with two-component Gaussian Mixture Model fits overlaid for the light-lumen (light blue) and dark-lumen (dark blue) subpopulations. **(D and E)** Fractional histogram and cumulative distribution of distances from light- (light blue) and dark-lumen (dark blue) vesicles to the plasma membrane. Vertical dotted lines in **(E)** indicate interquartile ranges for each population. **(F and G)** Fractional histogram and cumulative distribution of distances from light- and dark-lumen vesicles to the nearest microtubule. Vertical dotted lines in **(G)** indicate interquartile ranges for each population.

We took two independent strategies to quantify the number and size of these free vesicles. First, we created a supervised U-net in Dragonfly and applied it to 30 randomly selected tomograms. After processing, 1,690 total vesicles were recovered, with 1,359 light-lumen vesicles (80%) and 331 dark-lumen vesicles (20%). Light-lumen vesicles had a mean diameter of 53 nm, while dark-lumen vesicles had a mean diameter of 102 nm (Fig. 3B). Second, we segmented a set of 107 of the highest-quality tomograms and subsequently processed them through the Surface Morphometrics analysis pipeline (Barad et al., 2023), which provided morphometric and spatial relationships among all segmented structures (e.g., PMs or MTs). Based purely on size, an unsupervised model resolved the single-membrane vesicles into a bimodal distribution largely representative of the light- and dark-lumen distribution of the Dragonfly dataset (Fig. 3C). Across the 107 tomograms, we observed a smaller diameter vesicle population (68%) centered at 53 ± 6 nm (n = 2,921) and a larger diameter vesicle population (32%) centered at 86 ± 28 nm (n = 1,076). The two analysis methods yield similar populations of light-lumen and dark-lumen vesicles, with similar diameters, further indicating that there are two populations of free vesicles in DRG axons.

The Surface Morphometrics analysis not only reveals vesicle morphology information (e.g., class diameters), but also their spatial relationships with other organelles and with varicosity boundaries (PMs). We investigated whether the light- and dark-lumen vesicles occupied distinct spatial regimes relative to both the PM and MTs (Fig. 3D-3G). We found that the median distance from the plasma membrane was 153 nm for light-lumen vesicles and 166 nm for dark-lumen vesicles (Fig. 3D and 3E). Given the wide distribution of distances for both vesicle types, there appears to be little evidence of vesicle docking to the PM, suggesting these vesicles are not involved in activity-dependent evoked release and are more likely engaged in transport and/or intracellular membrane recycling. To test whether these vesicles were engaged in transport along MTs, we calculated their median distance from MTs. We found that light-lumen vesicles and dark-lumen vesicles had median distances of 125 nm and 121 nm from MTs, respectively (Fig. 3F and 3G). The two populations did not differ significantly in their proximity to MTs (p = 0.42). Further, the distances from MTs for both vesicle types span a broad distribution, suggesting that the majority of the free vesicles are not significantly engaged in MT-based transport.

### Endosomal and multilamellar compartments are morphologically distinct

In addition to free vesicles, we observed endosomal and multilamellar compartments that were significantly larger in size than the free vesicles. These compartments likely represent varying maturational stages along the endolysosomal pathway. We categorized these compartments into three classes: endosomes, multivesicular bodies (MVBs), and multilamellar vesicles (MLVs) (Fig. 4A and 4F, S2-S4). Across all tomograms, endosomes, MVBs, and MLVs were identified in 53%, 52%, and 33%, respectively (Table 1).

**Figure 4.**
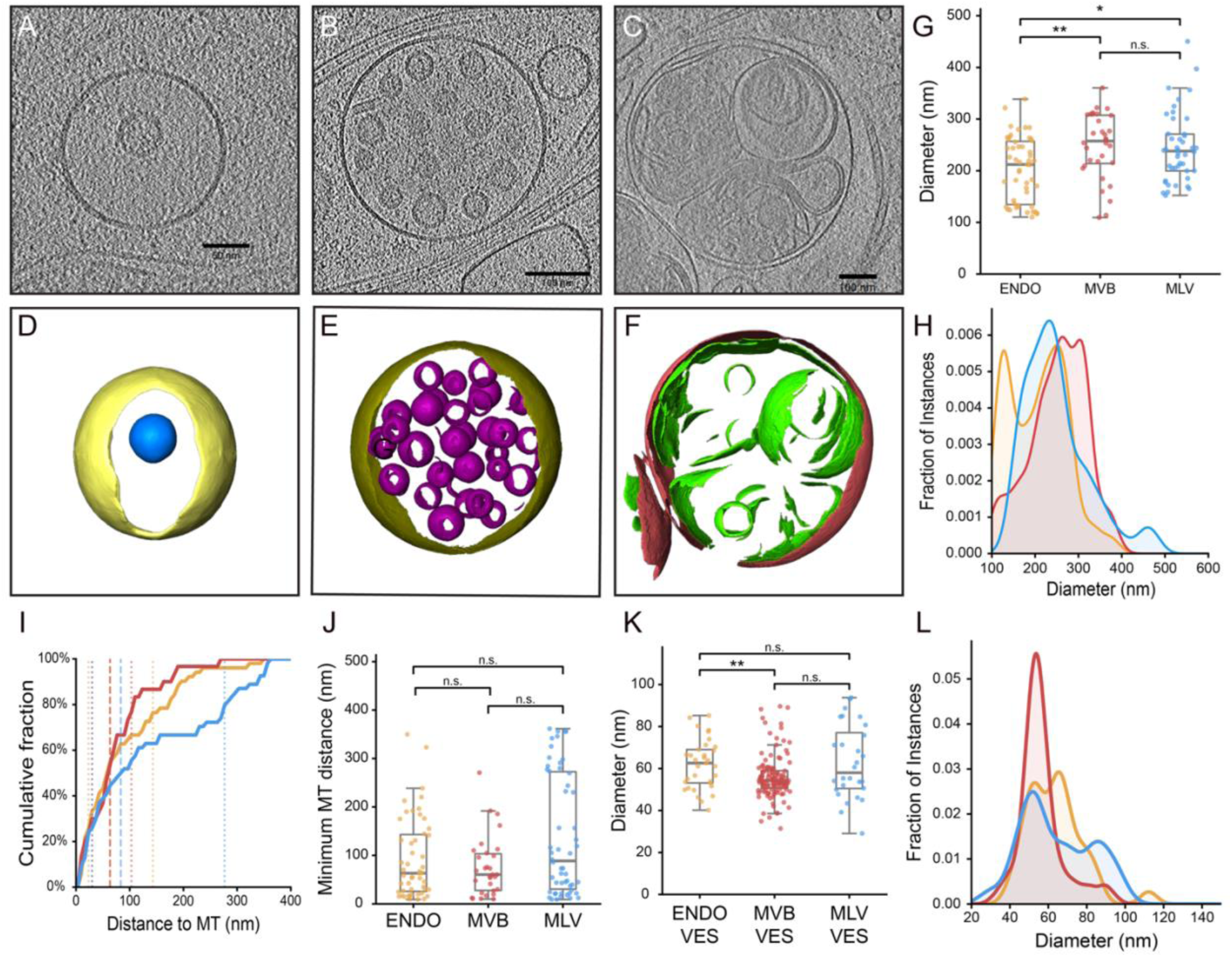
Size and spatial distributions of recycling and degradative compartments. (A-C) Slices from denoised tomograms as exemplars of endosomes, MVBs and MLVs, respectively. **(D-F)** Segmented volumes of the tomographic slices shown in A-C; endosome in yellow with its internal vesicle in blue, MVB membrane in brown with its internal vesicles in purple, and MLV membrane in brass with its vesicular and luminal membranes in green. **(G)** Per-instance diameters of endosomes (ENDO; 212.0 ± 62.4 nm, n = 51), MVBs (257.4 ± 63.4 nm, n = 32), and MLVs (238.0 ± 67.4 nm, n = 44). **, P < 0.01 for ENDO versus MVB; *, P < 0.05 for ENDO versus MLV; ns, not significant for MVB versus MLV; Mann-Whitney U test. **(H)** Fractions of instances distributions of diameters of endosomes, MVBs, and MLVs. MVB and MLV distributions appear unimodal while the endosomal distribution shows a bimodal character. **(I)** Cumulative distribution functions of minimum distance to the nearest microtubule for endosomes (orange), MVBs (red), and MLVs (blue). Vertical dashed lines indicate the 25th, 50th, and 75th percentiles for each class. **(J)** Minimum distance from each ENDO, MVB, and MLV compartment to the nearest MT. Median minimum distances were 63.6, 60.4, and 91.5 nm for ENDO, MVB, and MLV, respectively; ns, not significant between any class pair; Mann-Whitney U test. (**K)** Per-instance diameters of internal vesicles of endosomes (ENDO_VES; 62.6 ± 11.8 nm, n = 33), MVBs (MVB_VES; 54.2 ± 10.4 nm, n = 123), and MLV_VES (58.0 ± 17.6 nm, n = 28). **, P < 0.01 for ENDO_VES versus MVB_VES; ns, not significant for all other comparisons; Mann-Whitney U test. **(L)** Fraction of instances distributions of equivalent-sphere diameters of internal vesicles of the three compartment classes.

We quantitatively assessed the morphology of each of these classes using the Surface Morphometrics dataset, starting with the diameters of these compartments (Fig. 4G). We found that endosomes were significantly smaller in diameter than both MVBs and MLVs (Fig. 4G). We also analyzed the diameter distribution in each of these classes (Fig. 4H). We found that the distributions in the MVB and MLV classes were broad and unimodal, mirroring the size relationship of the diameter quantifications (Fig. 4G). The density distribution of the endosomal class was similarly broad but appeared bimodal, with peaks at approximately 125 nm and 240 nm (Fig. 4H). These distinct size profiles of endosomes suggest progressive remodeling across the endosomal pathway. Overall, the median diameter of these compartments ranged from 212 to 257 nm. What dictates their sizes is not clear but might be constrained by the significantly reduced internal diameter of the axon through which these vesicles will ultimately be transported. It is equally plausible that their diameters are constrained by the molecular machinery that regulates their biogenesis.

We further observed that these three multi-membrane compartments occupied distinct spatial regions relative to MTs. MLVs extended across the widest range of distances from the MTs with an upper quartile of 275 nm, whereas the upper quartiles for endosomes and MVBs were 147 nm and 107 nm, respectively (Fig. 4I and 4J). However, the medians of the minimum distances for the three are quite similar (endosomes: 64 nm, MVBs: 60 nm, MLVs: 92 nm) and not significantly different, in part due to the variability in the distances. Finally, we assessed the association of each organelle to MTs by placing a 15 nm edge-to-edge cutoff as a conservative proxy for organelles potentially undergoing MT-based transport. Electron microscopy and structural studies show that vesicles are linked to microtubules by motor complexes that position cargo membranes tens of nm from the microtubule surface (Kerssemakers et al., 2006; Urnavicius et al., 2018; Vale and Milligan, 2000). Using this metric, 28%, 27%, and 24% of endosomes, MVBs, and MLVs, respectively, were associated with MTs. The similar contact fractions suggest the three organelle classes are transported along MTs at comparable proportions. The absence of significant differences in MT proximity between classes suggests that MT engagement is not a distinguishing feature along the endolysosomal maturation axis within DRG axon varicosities.

### Internal contents differ by parent recycling organelle

We next examined the morphological characteristics of the internal content of endosomes, MVBs, and MLVs. Each of these organelle classes housed luminal membranes that appeared as either spherical vesicles in all three organelles or fragmented stacks or sheets in MLVs (Fig. 4A-4F, S2, S3, S4). We first quantified the diameters of the internal vesicles. MVB internal vesicles were significantly smaller than endosomal vesicles, whereas endosomal and MLV-associated vesicles were not significantly different from one another (Fig. 4K). These data suggest that distinct mechanisms may govern vesicle formation in MVBs compared to the endosomes and MLVs.

We also observed similarities and distinctions in the morphology of the material inside endosomes, MVBs, and MLVs. This was anticipated based on each organelle’s distinct role in membrane recycling. Endosomes contained fewer vesicles than MVBs, and the internal vesicle diameter of the latter was smaller, as noted above (Fig. 4K and 4L). Another distinguishing feature in MVBs was brick-shaped material in a subset of MVBs (Fig. S3H-S3I). MLVs discriminate themselves from endosomes and MVBs with the presence of possible degradative internal membranes and irregular stacks and sheets of membrane (Fig. 4F and S4). The differences in internal membranes across compartments likely reflect different stages along the endolysosomal pathway, with some at the late endosome or multivesicular body stage and others at the endolysosome or autolysosome stage (Klumperman and Raposo, 2014).

### Phagophores span a heterogeneous ultrastructural continuum

Phagophores are morphologically identifiable double-membrane structures representing the early stage of autophagosome formation. In cross-section, the tips of the inner and outer membranes are slightly enlarged, which ultimately fuse together to form the mature autophagosome (Bieber et al., 2022). We observed prototypical examples of phagophores in DRG varicosities (Figs. 5A-D), consistent with previous live-cell imaging of autophagosome formation in DRG axons (Maday et al., 2012; Stavoe et al., 2019). We identified phagophores at various stages of maturation in 10 of the 192 tomograms (Fig. S6 and Table 1). We also observed intact vesicles engulfed in the phagophore bowls (Fig. 5A-D). We also noted the electron lucent appearance in the luminal space between the inner and outer membranes as opposed to the particulate-like appearance of the cytosolic material being encapsulated (Fig. 5A and 5B). Phagophore membrane expansion requires a lipid supply from neighboring membranes, typically the ER (Maday and Holzbaur, 2014). We observed the intimate 3D relationship between the inner and outer phagophore membranes and the ER (Movie 4). Note how the ER appears to extend into the opening of the maturing phagophore, suggesting that, in this case, lipids are being supplied by the ER.

**Figure 5.**
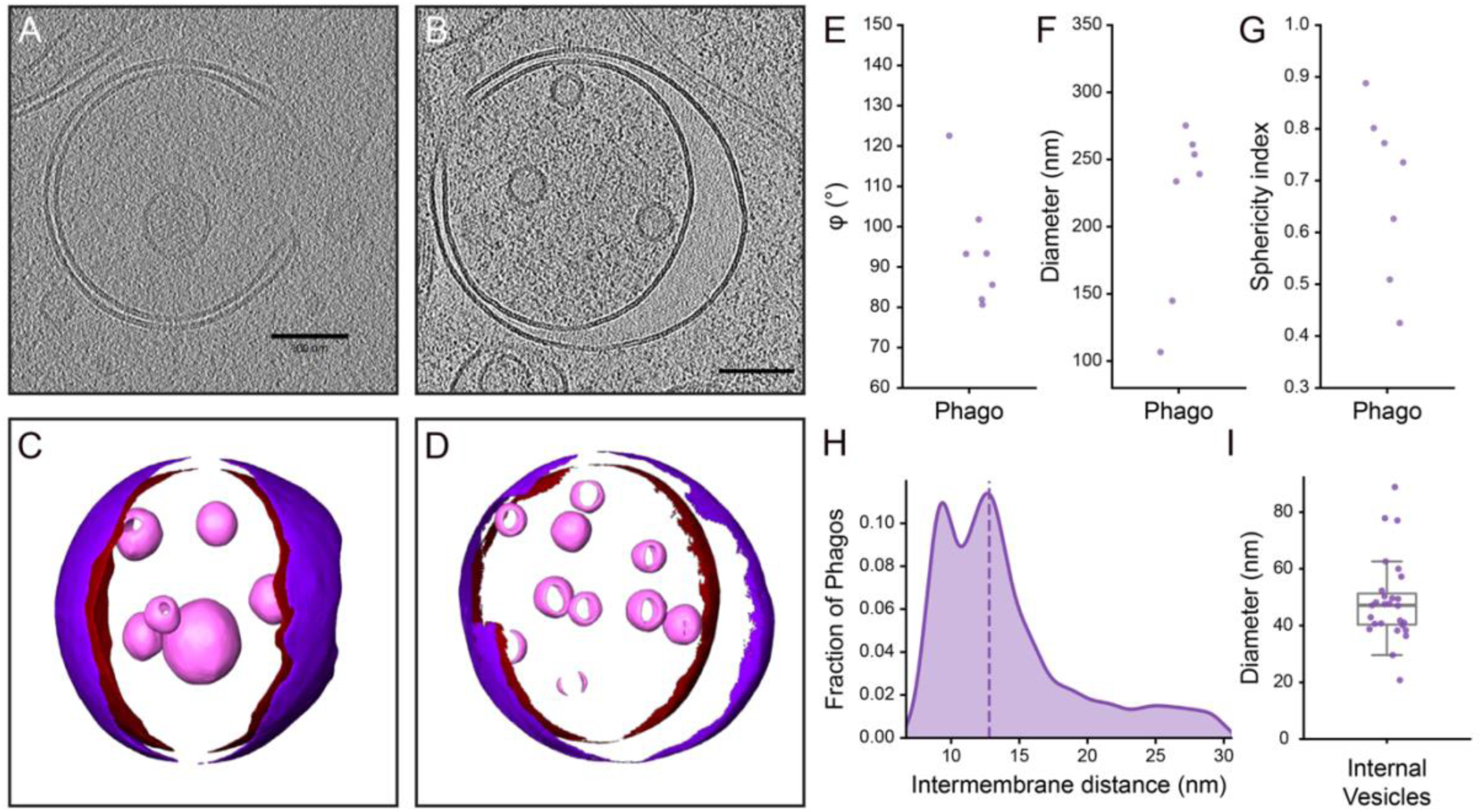
Phagophore ultrastructure and maturation states. (A and. **B)** Representative tomogram slices and **(C and D)** 3D segmentations of two phagophores at different maturation stages. Quantified geometric parameters for seven phagophores across tomograms. **(E)** Plot of the rim opening angle φ (degrees), calculated following Bieber et al. (2022) curvature-based approach, ranging from 80.7° to 122.5°. **(F)** Plot of the equivalent diameter (nm) derived from least-squares ellipsoid fit to inner membrane vertices, ranging from 106.6 to 275.2 nm. **(G)** Plot of the sphericity index (dimensionless, 0 = planar, 1 = perfect sphere), ranging from 0.43 to 0.89. Individual phagophores shown as scatter points. **(H)** Aggregate intermembrane distance distribution across phagophores with independently segmented inner and outer membranes. Dashed line indicates the median IMD of 12.8 nm. **(I)** Size distribution of internal vesicular cargo across five phagophores containing internal vesicles. The median diameter is 47.2 nm with a range from 20.8 to 88.9 nm.

Autophagosome formation involves progressive membrane expansion from an initiation site, cargo engulfment, and eventual closure to form a double-membrane autophagosome (Melia et al., 2020). Using data from the Surface Morphometrics pipeline, we calculated several morphometric parameters for seven phagophores. First, the rim-opening angles of the phagophores ranged from 80.7° to 122.5°, with a mean of 94.2° (Fig. 5E). These angles represent the steps of maturation as the tips of the phagophore close following a circular trajectory (Bieber et al., 2022). Second, we calculated equivalent diameters of the inner membrane of phagophores, which ranged from 106.6 to 275.2 nm (Fig. 5F). Third, the calculated sphericity of phagophores also suggests distinct developmental stages, with observed indices ranging from 0.43 to 0.89 (Fig. 5G). Lower index phagophores correspond to a more cup-shaped phagophore, whereas higher index phagophores correspond to a near-mature spherical autophagosome. Fourth, all seven phagophores were captured with sufficient clarity to independently segment their inner and outer membranes for an assessment of the intermembrane distance (Fig. 5H). The relatively narrow intermembrane space was variable along the curvature of the phagophore, with intermembrane distances ranging from 2.0 to 30.0 nm. These distances are consistent with previous reports of phagophore structure determined by cryo-ET in yeast (Bieber et al., 2022). Finally, we assessed the morphology of the internal vesicles captured by the maturing phagophores. Five of the seven phagophores contained internal vesicles. These 27 vesicles ranged in diameter from 20.8 to 88.9 nm, with a median diameter of 47.2 nm (Fig. 5I). These vesicles are likely cargo destined for autophagic degradation. None of the phagophores appeared to be associated with mitochondria or other identifiable organelles, suggesting they can be classified as part of the nonselective macroautophagy pathway.

### ER is a morphologically heterogeneous and ubiquitous compartment

The ER is an abundant organelle network that contacts and regulates most other organelle classes through membrane contact sites, which functionally link Ca²⁺ regulation, lipid transfer, and organelle positioning (Schrod et al., 2018; Foster et al., 2021). By visual inspection, the ER was present in 191 of 192 of the tomograms (Table 1). The morphology of the ER varied significantly, ranging from more tubular to more sheet-like (Fig. 1E, 1F, 6A, 6B). These two types of ER have been described previously (Pendin et al., 2011; Shibata et al., 2009). We calculated the curvedness and lumen width of the sheet-like and tubular ER (Fig. 6C and 6D). Tubular ER had a slightly higher median curvedness (0.018 nm⁻¹) than sheet-like ER (0.016 nm⁻¹). Lumen width also differed slightly, with median values of 53.3 nm for the tubular ER and 49.5 nm for the sheet-like ER. The fraction of ER surface area within 15 nm of an MT was not significantly different between both tubular and sheet-like ER (median 1.7% and 1.2%, respectively; p = 0.23), suggesting that both ER types interact with MTs similarly.

**Figure 6.**
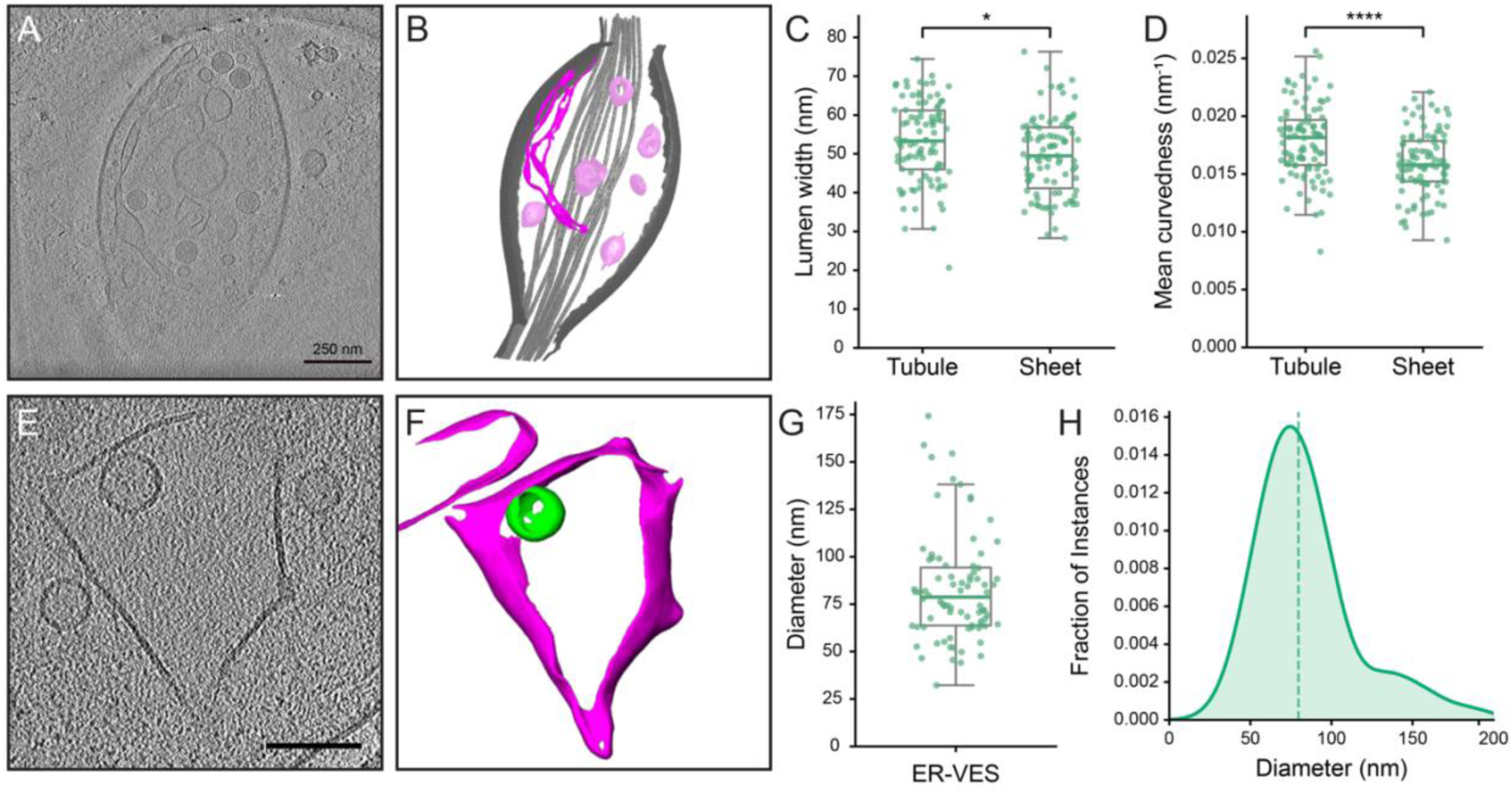
ER ultrastructure is heterogeneous and resolves into two broad classes. **(A)** Off-axis tomographic slice of a varicosity as an example of ER. Scale bar is 250 nm. **(B)** Segmentation showing a 3D view of the tomographic reconstruction in (A) to highlight the sheet (magenta) and tubular (pink) classes of ER. The MTs and PM are in greyscale; other organelles have been stripped away. **(C)** Plot of the luminal width of the tubular and sheet-like ER structures. *, P < 0.05; Mann-Whitney U test. **(D)** Plot of the mean curvedness of the membranes of the tubular and sheet-like ER structures. ****, P < 0.0001; Mann-Whitney U test. **(E)** Sheet-like ER from a tomogram distinct from that shown in (A) highlighting an internal vesicle. Scale bar is 100 nm. **(F)** The tomographic reconstruction in (E) is shown as a 3D segmentation to highlight that the vesicle (in green) is surrounded by ER membrane (in magenta). **(G)** Plot of the median diameter of vesicles found within the ER (n = 81 vesicles, 42 tomograms; per-tomogram median = 82.1 nm). **(H)** Fraction of instances distribution of the diameter of vesicles identified within ER. The distribution is largely unimodal with an elongated tail. The dashed line indicates the median diameter of 79.6 nm.

In many sheet-like ER compartments (36%), we detected discrete vesicles (Figs. 1H, 6E, 6F S7). To our knowledge, ER vesicles have not been described previously by cryo-ET in any cell type. These ER-associated vesicles had a median diameter of 78.7 nm (n = 80) (Fig. 6G). The distribution of ER-vesicle size was largely unimodal, with a tail at larger diameters (Fig. 6H). The functional significance of ER-associated vesicles is unknown, but their consistent presence in DRG neuron varicosities suggests a potential role in ER membrane remodeling.

A final feature of the ER we observed in a small subset of tomograms was a beads-on-a-string appearance (Fig. S6G and S6H). While a small sample, the beads had a diameter of 42 ± 11 nm (n=16) and the strings had an average diameter of 14 ± 2 nm (n=15). Such structures were also evident in DRG cryo-tomograms in a previous study of DRG axons in which they were termed “beaded-ER” (Foster et al., 2021). Similar structures were also evident in tomograms of cryo-preserved axons from human brain organoids (Hoffmann et al., 2021). This morphology is grossly similar to that observed in mitochondria subjected to forces related to osmotic pressure, membrane packing, and/or membrane tension (Sturm et al., 2025) and has been noted in previous EM studies focused on neuronal ER (Terasaki, 2018).

### Mitochondria are highly variable in dimensions and remodel at varicosity boundaries

One organelle that could be unequivocally identified for classification was the mitochondrion. We identified at least one mitochondrion with the typical double membrane structure and unique cristae morphology in 53% of the tomograms (Table 1, Fig. 1C-E). The morphology of the mitochondria was heterogeneous in size and diameter, and the density of cristae per mitochondrion also covered a broad range (Fig. 7A-C and S7). Another feature in many of the mitochondria was electron-dense precipitates (Fig. 6A - 6C). The precipitates were visualized in most mitochondria (83%; 85/102), but their number and density varied within individual mitochondria. These precipitates are reported to be calcium phosphate and appear to be functionally connected to the role mitochondria play in calcium buffering. Whether these precipitates are physiological or pathological remains an area of discussion (Anderson et al., 2026; Rose et al., 2025; Lycas et al., 2024; Wolf et al., 2017).

**Figure 7.**
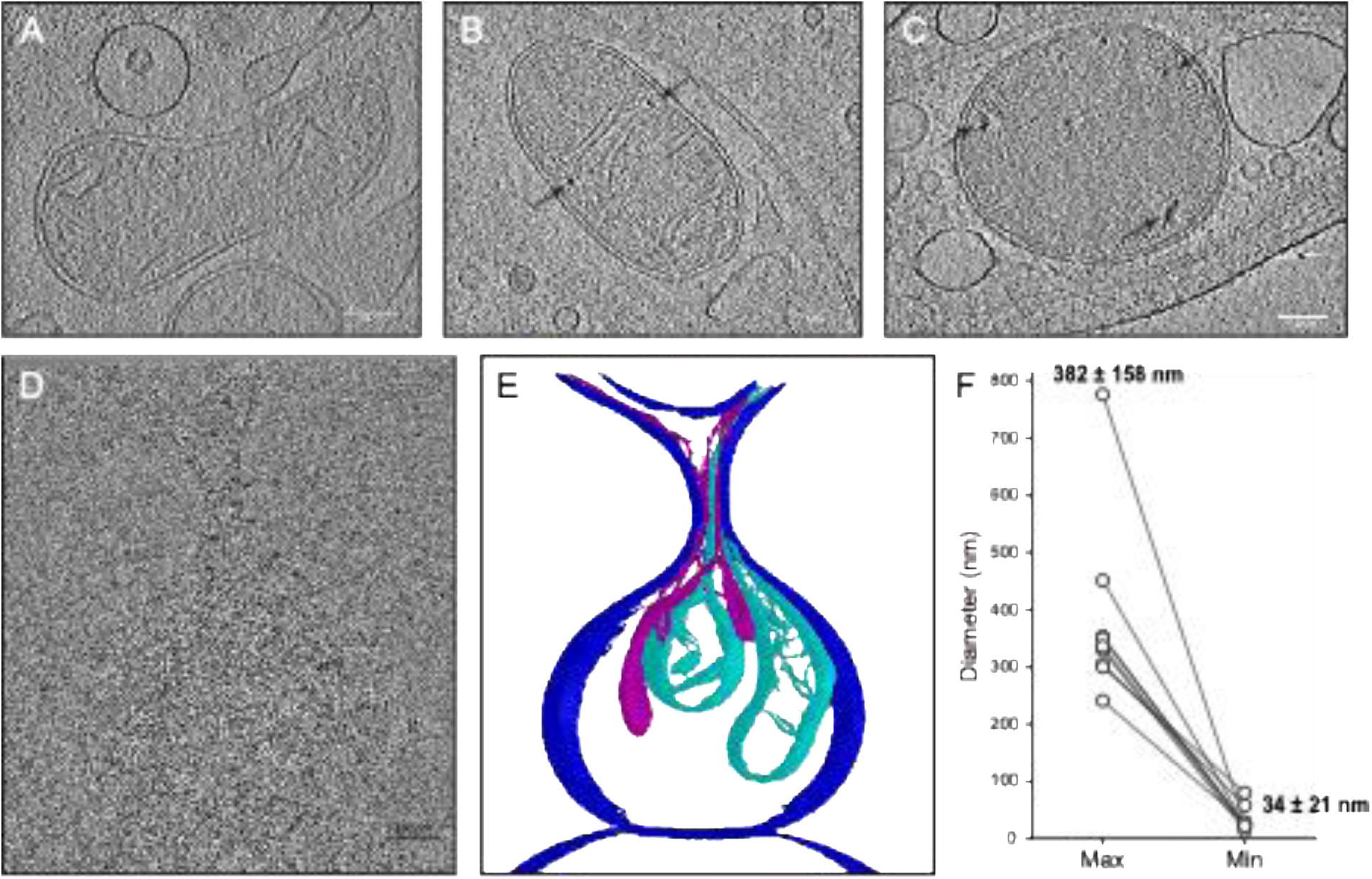
Mitochondria are heterogenous and remodel at axon/varicosity boundaries. **(A)** Slice through a tomogram of a varicosity showing a typical mitochondria with well-developed cristae. **(B and C)** Slices from two tomograms showing distinct mitochondria that exhibit electron-dense precipitates (highlighted with arrows). Scale bars are 100 nm. **(D)** Slice through a tomographic reconstruction including the axon and axonal/varicosity boundary at the top of the image. The outer and inner membranes of the two mitochondrion can be seen thinning as they exit the varicosity into the axon at the top right of the image (arrowheads). Scale bar is 250 nm. **(E)** A 3D segmentation of the tomogram shown in (A) that highlights the remodeling of the mitochondrial membranes (light blue) and a portion of the ER encircling one of the mitochondria (magenta). The plasma membrane is colored in dark blue. **(F)** Pair-wise plot of the difference in diameter of a collection of mitochondria caught at the axonal/varicosity transition. The average maximum diameter of mitochondria in varicosities is 382 ± 158 nm while the average minimum diameter of the thinnest portion of the extruded mitochondria is 34 ± 21 nm, n = 9.

Mitochondrial morphology in the DRG axons was notable at boundaries where the varicosity narrows in the transition into the axon shaft. We observed that a mitochondrion can occupy both distinct geometric regions: one part in the wider varicosity and another in the narrower shaft (Fig. 7D-7E). To accommodate these different boundary constraints, a significant remodeling of the mitochondrial membrane systems was required. We observed a portion of the ER surrounding the mitochondrion being extruded into the axon shaft (Fig. 7E). The outer mitochondrial membrane can be seen constricted in accordance with the narrowing of shafts as if being extruded, perhaps by motor-induced pulling into the axon (Fig. 7D). Concordantly, the inner mitochondrial membrane likely remodels, including dissolution of the cristae, and removing mitochondrial matrix from the constricted regions. The thinning and extrusion of the mitochondrion is dramatic and, in some examples, is narrowed to a diameter similar to that of ER tubules and MTs in the axons (Movie 5 and Fig. S8). As a metric for this thinning, the diameter of the mitochondria in each tomogram was compared within the varicosity to the thinnest diameter within the axon (Fig. 7F). The average diameters of mitochondria decreased from 382 nm to 34 nm. Several of the thinnest examples appeared to reach an asymptotic value of ∼24 nm, suggesting the minimum diameter mitochondria can attain. We suggest that remodeling of mitochondrial is related to their transport, constrained by the thin diameter of the axonal shaft.

## DISCUSSION

This study establishes a quantitative ultrastructural framework for understanding how membrane-bound organelles are organized within the physically constrained environment of developing axons. By combining cryo-electron tomography with quantitative morphometric analyses, we identify organizational features that span multiple trafficking pathways, including distinct vesicle populations, endolysosomal and autophagic intermediates, ER-associated vesicles, and dramatic mitochondrial remodeling events. Collectively, these observations suggest that spatial confinement imposed by axonal geometry is a major determinant of organelle architecture and membrane remodeling within the axonal compartment. More broadly, our findings indicate that organelle remodeling is not solely a consequence of maturation and trafficking, but also an adaptation to the physical constraints imposed by the heterogeneous geometry of axons and varicosities. The developmental stage of the DRG neurons is an important consideration when interpreting these observations. DRGs at 48 hr in culture are in an active growth phase characterized by robust axonal extension (Malin et al., 2007). We specifically selected this timepoint to quantify organelle structure and distribution during a period of extensive membrane remodeling associated with axonal outgrowth. In addition, DRG neurons at this stage exhibit little spontaneous activity (Black et al., 2018), and the composition and distribution of organelles should be interpreted within this developmental context. In this regard, the present manuscript complements previous cryo-electron tomography investigations of DRG axons that focused primarily on cytoskeletal organization (Foster et al., 2021).

Using two different segmentation strategies, we found the single-membrane vesicles to segregate into two distinct populations by lumen contrast and diameter. Using both techniques, the dark-lumen population was found to be significantly larger with a broader distribution than the light-lumen population and their sizes closely paralleled previously reported data (Foster et al., 2021). The tight distribution of the light-lumen vesicles indicates their size is under more stringent control than dark-lumen vesicles. Both populations exhibited similar spatial distributions relative to the PM and MTs and both showed more proximity to MTs than the PM. This argues that at this developmental stage of the cultured DRG neurons, the two vesicle populations follow similar trafficking routes and are not distinguished by different cytoskeletal transport processes. Single-membrane vesicles are known to serve multiple functions in axons. The more abundant light-lumen vesicles likely correspond to synaptic vesicles, synaptic vesicle precursors, or early endosomes based on similarities in diameter (Tao et al., 2025; De Pace et al., 2020; Schrod et al., 2018; Wang et al., 2016; Watson et al., 2023). The more heterogeneous and larger dark-lumen population likely corresponds to dense core vesicles (DCVs) that carry neuropeptides or neurotrophins (Sorra et al., 2006; Tao et al., 2018; Zhai et al., 2001; Bost et al., 2017; Xia et al., 2009).

The capability of cryo-ET to capture static snapshots of cellular activity led to observations of a continuum of endosomal and multilamellar compartments spanning multiple ultrastructural states. We detected MLVs in just over 30% of tomograms, indicating that although they are less prevalent than endosomes and MVBs, they remain a common feature of DRG axons and appear to represent a specialized late stage within the endolysosomal maturation pathway. Further, we found the endosomes, MVBs, and MLVs to have overlapping but non-identical diameter distributions, consistent with their roles across different stages of maturation through the endosomal pathway (Jordens et al., 2001; Mottola, 2014). Interestingly, despite different internal contents, the diameter of the limiting membranes of all 3 types of recycling vesicles appeared to reach a similar maximum size. It is possible that there are mechanisms in place to constrain the size of organelles for efficient trafficking along MTs through the significantly thinner and crowded axonal shaft. Spatially, the three compartments showed overlapping MT-distance distributions, further supporting that these organelles do not show differential probability of transport on MTs. However, our tomograms did not have the contrast or resolution to provide evidence of direct protein-mediated connections with the MTs.

The luminal content of endosomes, MVBs and MLVs varied significantly. MLVs exhibited broad, heavy-tailed internal vesicle diameter distributions, consistent with vesicles whose size emerges from graded membrane reorganization rather than templated budding. In this context, vesicle diameter is influenced by cargo engulfment, fusion history, and lipid-driven multilamellar assembly rather than a single curvature-imposing scaffold, producing substantial size heterogeneity (Mizushima et al., 2010; Nixon, 2013). This contrasts with ESCRT-mediated intraluminal vesicle formation in MVBs, which enforces tight size regulation and yields a narrow diameter distribution consistent with templated budding under strong curvature control (Klumperman and Raposo, 2014; Von Bartheld and Altick, 2011; Wollert and Hurley, 2010). Endosomal vesicles displayed intermediate variability, suggesting a mix of molecular mechanisms that generate internal endosomal vesicles. Although the origin of the endosomal vesicles cannot be determined from morphology alone, they may arise through ESCRT-mediated inward budding of the limiting membrane or through fusion with vesicle-containing endosomal intermediates. Their presence supports the view that endosomal maturation occurs along a continuum of structural states rather than through discrete transitions between organelle classes.

ER is a ubiquitous axonal compartment and exhibits a variety of sheet-like and tubular structures. Functionally, axonal ER serves as a platform for calcium storage and signaling, lipid synthesis and transfer, and organelle contact sites (notably ER–mitochondria and ER–endolysosomal contacts), all of which are critical for axonal homeostasis and long-range signaling (Terasaki, 2018; Kuijpers et al., 2021). The ER has been identified in numerous studies of cryo-preserved neurons (Foster et al., 2021; Schrod et al., 2018; Hoffmann et al., 2021; Nedozralova et al., 2022; Fischer et al., 2018) and takes on a variety of morphological shapes. Our data align with previous studies that broadly categorized ER into two types, sheet-like and tubular (Terasaki, 2018). From morphology alone, it is not possible to speculate what the functional roles these two distinct forms of ER might provide but is a point of interest for future studies.

A unique observation made in our tomograms is the presence of vesicles within the lumen of the ER in over a third of our tomograms. These vesicles appear to be entirely enclosed within the ER lumen. To our knowledge, vesicles within the ER have not been described in prior cryo-ET studies of ER. Their composition and functional roles are unknown but there are several intriguing possibilities. These vesicles may represent COPII-independent intra-ER transport intermediates (Gaynor and Emr, 1997; Peotter et al., 2019). Alternately, they may represent early ER-phagy events in which ER tubules are being packaged into an isolation structure (Grumati et al., 2018; Hoyer et al., 2024), or they may represent lipid droplet precursor intermediates (Wilfling et al., 2014). Importantly, these ER-vesicles will require further study to dissect their composition and potential function.

Varicosities are sites of concentrated endolysosomal and autophagic traffic, where autophagosomes that formed distally mature through fusion with late endosomes and lysosomes (Maday and Holzbaur, 2014; Stavoe et al., 2019; Cheng et al., 2015). We identified phagophore-like structures in a minority of tomograms. Given that autophagosome biogenesis occurs at a rate of approximately once every 2 min in DRG neurons cultured from young adult mice (Stavoe et al., 2019), it is unsurprising that we captured a limited number of phagophores in our tomography datasets. The phagophore interluminal space was electron lucent and distinct from the granular appearance of the material engulfed within the internal membrane of phagophores. Because the interluminal space is topologically equivalent to the inside of an organelle (e.g., from the fusion of ATG9 vesicles or omegasomes on the ER; (Hurley, 2026)) its electron lucent appearance is expected and appears largely devoid of proteinaceous material (Li et al., 2023; Bieber et al., 2022). In addition, the presence of internal vesicular cargo and general cytoplasmic constituents in the majority of observed phagophores suggests active sequestration of cytoplasmic material consistent with constitutive axonal autophagy (Maday and Holzbaur, 2014; Stavoe et al., 2019; Yang and Klionsky, 2010). Using live-cell imaging with fluorescently tagged organelles, a previous study determined that the donor membrane for the expansion of the phagophore membrane in neurons appears to be largely from the ER (Maday and Holzbaur, 2014). While we had limited phagophore samples, our data directly observed a close spatial relationship between the ER and phagophore (Movie S2), consistent with the ER as the source for donating lipids to the growing phagophore in axons.

Mitochondria were the easiest organelle to unequivocally identify in our dataset. The vast majority of mitochondria (83%) exhibited internal calcium phosphate precipitates. Our data support the physiological importance of these precipitates in the normal function of buffering/storing calcium (Anderson et al., 2026; Wolf et al., 2017; Lycas et al., 2024). Future studies employing poisons of mitochondrial membrane potential (Wolf et al., 2017) would help confirm the reversible nature and physiological role of these intramitochondrial precipitates. Beyond their internal composition, however, mitochondria provided the most striking example of the structural adaptations required for transport through the spatially constrained environment of axons. A striking feature of our tomographic data was the diameter of mitochondria within varicosities relative to the significantly thinner axons that the mitochondria must traverse for transport. This physical barrier was noted previously for mitochondria in hippocampal neurons also examined by cryo-ET (Fischer et al., 2018). Mitochondrial transport in DRG neurons cultured for 48 hr (as in our study) showed that the proportion of mobile vs immobile mitochondria was approximately 25%/75% in live cell imaging experiments (Moughamian et al., 2013). It is intriguing to conjecture that the large fraction of non-motile mitochondria is in part a consequence of this physical barrier to transport. At varicosity boundaries, we observed mitochondria to constrict the OMM to fit into the narrowing axon diameter. The IMM remained intact in the varicosity and closely followed the constriction of the OMM. However, the cristae disappeared as the OMM and IMM narrowed. We presume this is a necessary and active remodeling process to permit the mitochondrion to narrow sufficiently to navigate the limited axonal space. Previous studies found that in the process of mitochondrial fission, the outer and inner membranes are thinned to a diameter of 15-50 nm with a disappearance of the internal cristae structure before the membranes undergo fission (Mageswaran et al., 2023; Kalia et al., 2018). An interesting possibility is that some of the molecular machinery necessary to produce mitochondrial fission (Kalia et al., 2018) is utilized in the remodeling noted in our tomograms. Under normal conditions, the OMM and IMM are connected at cristae junctions via the MICOS (mitochondrial contact site and cristae organizing system) complex (Mageswaran et al., 2023; Kalia et al., 2018; Pfanner et al., 2019). Presumably, the constriction of the OMM would disrupt the MICOS contacts to permit the IMM to retain its integrity as it narrows along with the OMM while the cristae disappear. A likely consequence of this remodeling is that dissolution of the cristae-rich mitochondrial domain would impair oxidative phosphorylation and reduce ATP production (Cogliati et al., 2013). Regardless of the underlying mechanisms involved, our observations demonstrate that substantial mitochondrial remodeling is required to accommodate the geometric constraints imposed by the axonal compartment. More broadly, mitochondria provide perhaps the clearest example of how spatial confinement shapes organelle architecture and function within neurons, reinforcing the idea that adaptation to axonal geometry is a common feature of multiple membrane trafficking pathways.

### Outlook

The present work establishes a quantitative framework for understanding how organelles are organized within the physically constrained environment of developing axons and highlights how cryo-ET combined with emerging morphometrics approaches can provide detailed quantitative description of cellular architecture. An important challenge going forward will be to connect the diverse morphologies captured here, including the continuum of endolysosomal intermediates, the distinct vesicle populations, and vesicles observed within the ER lumen to specific molecular mechanisms and trafficking pathways. In parallel, our observations suggest that the narrow and heterogeneous geometry of axons is an important determinant of organelle architecture, membrane remodeling, and transport. Future studies will be required to determine how spatial confinement influences organelle function and how these constraints are integrated with the molecular machinery that governs trafficking and membrane dynamics. Extending these analyses across developmental stages, physiological states, and aging will be essential for determining how these organizational principles are remodeled over time. Because DRG neurons can be isolated from animals across the lifespan, they provide an ideal model for examining age-related changes in axonal organelle organization and trafficking, a major focus of our ongoing work. Collectively, our findings support a model in which adaptation to axonal geometry is a common feature of multiple membrane trafficking pathways and establish spatial confinement as a fundamental organizing principle of axonal cell biology.

## MATERIALS AND METHODS

### Culture of DRGs for EM

DRGs were isolated from 3-month-old mice exactly as previously described (Waxham et al., 2026). All procedures associated with the mice were performed in accordance with guidelines established by the Animal Care committee at UTHealth Houston. Briefly, mice were euthanized by decapitation after terminal anesthesia, and the spinal cord was removed. The ganglia from cortical to thoracic regions were dissected and digested with papain for 20 min at 37°C with occasional mixing. DRGs were collected by centrifugation and then treated with collagenase/dispase for 20 min at 37°C with occasional mixing. The ganglia and partially dissociated cells were collected by centrifugation, and the pellet was triturated with a fire-polished glass pipet that was previously coated with complete F-12 medium containing serum. Dissociated cells were collected by centrifugation through at 20% Percoll cushion and resuspended in complete F-12 media by gentle pipetting. Cell counts were made; the cells were collected again by centrifugation and finally resuspended targeting a volume where 5,000 live DRG neurons were plated on each EM grid. The previous day, gold Quantifoil SiO_2_ or carbon grids (2/2) were glow discharged and placed into PLL (100 mg/ml) overnight at 4°C in 35 mm Mat-tek dishes, 4 grids per dish. The grids were placed on the plastic surrounding the glass coverslip depression to facilitate later pickup with forceps. The grids were washed with sterile water, briefly air dried, and then coated with laminin (20 mg/ml) for 2h at 4°C and placed in complete F12 medium before DRG plating. For plating, the concentrated DRG cell solution was added directly to the top of the EM grids incubating in complete F12 medium, that were then gently moved to the incubator and stored for 2d at 37°C/5%CO_2_. For the data presented in this manuscript, DRGs were isolated and cryopreserved from three separate 3-month-old mice.

### Cryopreservation of DRGs on EM grids

Vitrification of DRGs was accomplished using a manual blotter at room temperature setup adjacent to the cell incubator, minimizing the time between removal of grids from the 37°C environment and vitrification exactly as detailed (Waxham et al., 2026). Each grid was shuffled to the edge of depression of the Mat-tek dish and picked up by their edge with forceps attempting to minimally bend/disturb the grid surface. 4 μL of BSA coated 10 nm fiducial gold in F-12 media was added to the grid surface and the forceps were placed in the manual blotting apparatus. The grid was blotted from behind with 1M Whatman filter paper for approximately 3-5 sec and then plunged into liquefied ethane at liquid nitrogen temperature. Each grid in a dish was processed sequentially over an approximately 2 min period, attempting to minimize the time cells spent at reduced temperature before cryopreservation. The grids were stored until imaged in a liquid nitrogen Dewar.

### Acquisition of cryo-ET datasets

All data in this manuscript were collected on one of two cryo-electron microscopes operated by the Structural Biology Center at UTHealth-Houston. Most data were collected on a Titan Krios operated at 300 keV equipped with a Gatan energy filter and a K2 Summit camera. A comparative subset of data was collected on a Glacios operated at 200 keV equipped with a Selectris energy filter and a Falcon 4i camera. While contrast was somewhat improved from data collected on the Glacios, there were no differences noted in the structural information of the data between the two machines so all the data for qualitative and quantitative analyses were combined. Data acquisition was accomplished on EM grids clipped in cartridge/C-clips using SerialEM of TOMO5 software. Following acquisition of atlases (175x) areas of appropriate thickness ice with well isolated DRG axons were targeted for collection of montages (8500x). At this magnification axons and varicosities with organelles were discernable and sites were targeted for data acquisition tracking along the axons. Tilt-series were acquired at each position from -51 to + 51° at 3° tilt increments. Data for each tilt were collected in dose-fractionation mode at a pixel size of 5.4 Å/pixel (23kx magnification). Total dose for each tilt-series was between ∼120-180 e^-^/Å^2^ and defocus was kept to 6-8 μm. Condenser and objective apertures were 70 μm and the energy filter was set to -20 eV in zero loss mode. All frames were acquired gain-corrected and stored in the .mrc file format at 16 bits. Data was acquired on at least two grids from each of the three mice.

### Data pre-processing and tomogram production

Custom written scripts were used to create a pipeline to process the raw movie frames into a tilt series. These and other associated scripts for data processing can be found on the Zenodo repository (10.5281/zenodo.16927937) and detailed methodology can be found in Waxham et al., (Waxham et al., 2026). Briefly, movie frames were corrected with alignframes (IMOD) and then the aligned frames were reassembled into a single tilt-series stack. The drift-corrected tilt-series were imported into BatchRunEtomo to produce 4x binned (21.6 Å/pixel) weighted back-projection tomograms. Manual screening eliminated poorly aligned or poorly targeted acquisitions. All tomograms passing this step were denoised by processing through IsoNet’s deconvolution program that improved contrast and signal-to-noise. For these datasets we found that changing the SNRFalloff to 0.6 and the DeconvStrength to 1.5 produced optimal denoising. These tomograms were used for subsequent downstream processing as well as for manual annotations of organelle numbers. For some purposes, 2x binned tomograms were produced (10.8 Å/pixel), returning to eTomo, adjusting parameters and running the data through the alignment and denoising steps recursively.

### Data processing for Surface-Morphometrics

Of the 192 reconstructed tomograms, 107 were deemed to be worthy of segmentation based on tomogram quality. First, membrane voxels were segmented through the deep-learning program MemBrain-Seg (Lamm et al., 2025) using its default parameters. Then, microtubule voxels were segmented using the deep-learning model TARDIS-MT (Kiewisz et al., 2025). The outputs of each were imported as separate label files into Amira (Thermo Fisher Scientific), where the files were manually segmented. A 16 class profile was established for each tomogram that included: outer mitochondrial membrane (OMM), inner mitochondrial membrane (IMM), plasma membrane (PM), microtubule (MT), endoplasmic reticulum (ER), ER-vesicles (ER_VES), multi-vesicular body (MVB), MVB-vesicles (MVB_VES), free vesicles (FREE_VES), multilamellar vesicle (MLV), MLV-vesicles (MLV_VES), phagophore (PHAG), PHAG-vesicle (PHAG_VES; this referred to the inner membrane of the phagophore), PHAG-internal vesicle (PHAG_INT_VES), endosome (ENDO) and ENDO-vesicle (ENDO_VES). Threshold-based selection and the 3D magic wand tool were used to assign membrane voxel regions into one of our semantic classes based on morphological criteria described in the results section. Completed segmentations were exported as MRC2014 files.

To quantify organelle size, shape and inter-organelle distances, the MRC2014 Amira output file was passed to the Surface Morphometrics workflow. For a thorough discussion of the details of this pipeline the reader is referred to (Barad et al., 2023) and the most recent versions of the program are available at https://github.com/GrotjahnLab/surface_morphometrics. This is a Python-scripted pipeline that takes parameters from a configuration.yml file for each dataset, called from three separate programs. First, the MRC2014 files with their associated semantic segmentations are converted to surface meshes output in the .vtp format. Second, each mesh is converted into a graph format (.gt) and curvature is calculated using pycurv (Salfer et al., 2020). Finally, distances and orientations are calculated from the .gt files and a final .csv file is created along with an updated .vtp file that can be imported. Finally, distances and orientations are calculated from the .gt files and a final .csv file is created along with an updated .vtp file. The .vtp files created at different steps can be used to visually assess (using Paraview; https://www.paraview.org/ ) whether issues might have arisen in processing data through the pipeline.

### Connected-component labeling and instance separation

The surface morphometrics pipeline produces per-class surface meshes that do not separate individual instances of an organelle. Each individual organelle instance was therefore isolated via a connected-component labeling algorithm applied to the pycurv triangle graph (.gt) files output by the pipeline. For each class, the .gt file was loaded using the graph-tool library (Peixoto, 2014)(v2.98). Connected components were labeled with the label_components function (directed = False). The component identifier (unique_component) was stored as a vertex property and exported to a per-class cc.csv file containing all per-triangle metrics from the pipeline. For quality control, components with fewer than 50 triangles or a total area below 150 nm^2^ were discarded as segmentation fragments.

### Microtubule geometry analysis

For each MT instance, the connected-component labeled mesh was converted to a point cloud of triangle centroids. A principal component analysis (PCA) axis was fitted to each instance to define the best-fit straight-line trajectory of each MT. The perpendicular distance from each triangle centroid to this axis was then computed, and the root-mean-square (RMS) of these distances was taken as the bending deviation for that MT instance. Straighter MTs yield smaller bending deviations, while more curved MTs yield larger values.

Apparent MT persistence length was then calculated as a comparative curvature-based metric. For each MT instance, the end-to-end span along the PCA axis was taken as the length L. The apparent persistence length was derived from the expression:

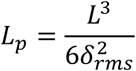

*δ_rms_* is the RMS bending deviation. This calculation follows the worm-like chain framework of Gittes et al. (Gittes et al., 1993; Wisanpitayakorn et al., 2022). MTs were restricted to lengths between 200 and 2,000 nm and excluded if their bending deviation was <1 nm before fitting the population distribution.

### Organelle size measurements

To measure the sizes of each organelle instance, a principal component analysis (PCA)-based semi-axis fitting procedure was applied to the vertex coordinates of each connected component identified via the procedure outlined above. For each component, the vertex coordinates were first centered around the component’s average position. The covariance matrix is decomposed by eigen analysis. Then, the point cloud is projected onto the three principal eigenvectors. Semi-axes are estimated to be half the peak-to-peak extent along each axis. The equivalent-sphere diameter is derived from the ellipsoid volume. A sphericity index was then calculated using the approach of (Cruz-Matías et al., 2019), where values closer to zero indicate a flat disk-like object and values closer to one indicate a more spherical object. Components were subject to minimum vertex thresholds that varied by class: 50 for free vesicles, 100 for outer compartment membranes (endosomes, MVBs, MLVs), and 20 for inner vesicle classes.

### Gaussian mixture model for single-membrane vesicles

Single-membrane vesicles were categorized into two subpopulations of a smaller and larger diameter using a two-component Gaussian Mixture Model (GMM) fitted to the diameter distribution. Prior to fitting, the following quality filters were applied: diameter within 20-500nm, principal axis ratio < 3.0, vertex count > 50, per-tomogram removal of top 1^st^ percentile by volume of the ellipsoid to suppress extreme outliers. The GMM was fitted with scikit-learn’s GaussianMixture (Pedregosa et al., 2011) with parameters (ncomponents = 2, random state = 0).

### Inter-Organelle distance analyses

Per-triangle distances to reference surfaces were calculated by the surface morphometrics pipeline as described in Barad et al. (Barad et al., 2023). We extracted a single minimum distance per organelle instance from the connected-component CSV files by identifying the closest triangle in that component to the reference surface. Distances to MTs were measured to the TARDIS-MT ribbon mesh, which approximates the MT centerline. The reported values therefore represent organelle surface-to-MT-axis distances.

Per-instance minimum distances were pooled across all tomograms and binned into fractional histograms of 60 equal-width bins over 0–400 nm. For the free vesicle distance analysis, the GMM population assignments were mapped to component IDs per tomogram via a lookup table, and histograms were computed separately for each population. MT-contact fractions were computed as the proportion of organelle instances with a minimum MT distance ≤27.5 nm that corresponding to approximately 15 nm edge-to-edge clearance and the ∼12.5 nm MT radius.

Area-weighted fractional histograms of minimum distance were computed per tomogram in 60 equal-width bins over 0–400 nm. For the free vesicle distance analysis, the GMM population assignments were mapped to component IDs per tomogram via a lookup table. Histograms were computed separately for each population, then normalized and averaged across tomograms. MT-contact fractions were computed as the proportion of tomograms containing at least one organelle instance with a minimum distance ≤15 nm.

### Phagophore morphology

The rim opening angle was calculated as the angle between the rim plane and tangential planes to the inner membrane close to the rim, ranging from 0° for a flat membrane disk to ∼180° just before closure. We read each phagophore inner membrane VTP file using the VTK XML PolyData reader and identified the topological rim using vtkFeatureEdges (BoundaryEdgesOn). Tomograms with fewer than 8 rim vertices were excluded. The rim plane normal vector was estimated as the smallest singular vector of the SVD (singular value decomposition) of the centered rim coordinates. Inner membrane vertices within 70 nm of the rim plane were divided into 10◦angular batches. A tangent plane was fitted to each batch by SVD. When the available rim arc exceeded 90°, the axis of the cone subtended by the tangential normals were used as a corrected rim normal. The rim opening angle is then computed from the relationship between the corrected rim orientation and the local tangential membrane orientations.

The intermembrane distances (IMD) were calculated using the refined bidirectional minimum distance algorithm of (Bieber et al., 2022). The mean IMD is computed by collecting refined nearest-neighbor distances between the paired membrane surfaces. Phagophores with extremely small intermembrane distances were excluded because such values indicate insufficient mesh separation between the two bilayers.

For each phagophore inner membrane, a minimum-volume enclosing ellipsoid (MVEE) was fitted using the iterative algorithm of (Todd and Yıldırım, 2007), (maximum iterations = 100, tolerance 10^−6^). Given semi-axes from the MVEE fit, the diameter and sphericity were computed as described above. Components with fewer than 20 vertices were excluded.

### Gaussian mixture model for ER sheet vs tubule classification

ER membrane was classified as tubular or sheet-like using fixed shape index thresholds applied to the per-triangle shape index values output by pycurv. The shape index is a dimensionless curvature descriptor ranging from −1 to +1, where values near +0.5 correspond to cylindrical (tubular) geometry and values near 0 correspond to planar (sheet-like) geometry. Triangles with shape index ≥ 0.5 were assigned to the tubular class; triangles with shape index ≤ 0.0 were assigned to the sheet-like class; intermediate values (0.0–0.5) were excluded. The fraction of ER membrane area belonging to each class was computed as the area-weighted proportion of triangles assigned to each category.

For each tomogram, the GMM was fitted with scikit-learn’s GaussianMixture (Pedregosa et al., 2018) with parameters (n_components = 2, random_state = 42, max_iter = 300). The component whose mean was nearest to +0.5 was assigned as the tubular class; the remaining component was assigned as the sheet-like class. The fraction of ER membrane area belonging to each class was computed as the area-weighted proportion of triangles assigned to each component.

### Data processing for Dragonfly

For quantifying the number of light and dark vesicles a trained neural network was first created in Dragonfly (Comet Industries) using 4 tomograms that were iteratively segmented and refined visually, classifying vesicles as either light or dark to establish a manually annotated training dataset. The training tomograms were selected to have a reasonable number of both light and dark free vesicles and were from the 4x binned, IsoNet denoised datasets. A network was then created using the Deep Learning Tool in Dragonfly. Parameters for U-net training were: batch size -32; Patch size – 128,128,3; Stride Ratio – 1; Data Augmentation Factor -5; Loss Function – Cross Entropy; and Optimization Algorithm – Adadelta. After asymptotic training, the network was applied to 30 randomly selected tomograms across the various datasets. Next, a cleaning step was applied removing islands composed of less that 500 connected voxels. Then, any vesicles that had been misclassified or perhaps classified with both light and dark portions, were manually cleaned. Vesicles were binary – dark or light. After applying a connected components filter to the data, a multi-ROI was produced for each of the dark and light vesicle populations. From there the built-in quantification functions of Dragonfly were employed. Individual vesicles were assessed for: Feret minimum, maximum and mean diameters, sphericity, and volume. The data in output .csv files to each tomogram was processed and calculations made using custom Python scripts.

## Supplemental material

Supplementary data are provided in Figures S1-S8 and Movies S1-S4.

## Data availability statement

Representative tomograms from each step of the processing pipeline, Python scripts for specific steps, and Dragonfly object files will be made available by the communicating authors.

## ACKNOWLEDGEMENTS

We gratefully acknowledge many helpful discussions with Dr. Matthew Swulius and members of the Stavoe and Waxham Labs. Dr. Jessica Heebner is acknowledged for expert help with Dragonfly network training and segmentation. This work was supported by NIH Grant R21AG090967 (to A.K.H.S. and M.N.W.). B.A.B. is supported by funding from the Collins Medical Trust. M.N.W. acknowledges the William Wheless III Professorship. The Glacios microscope was partially purchased through support from a grant from the NIH Office of the Director (1S10OD032204).

## Abbreviations

DRG: dorsal root ganglia
Cryo-ET: cryogenic electron tomography
SVD: singular value decomposition
GMM: Gaussian mixture model
MVEE: minimum-volume enclosing ellipsoid
IMD: intermembrane distances

## Supplemental figure and Video legends

Figure S1. **Gallery of tomographic slices of cytoskeletal elements in varicosities.** Panel A shows an example of long straight thick filaments (arrows) running through the center of the varicosity. MTs are indicated by the arrowheads. Panel B is a cartoon rendition of the cytoskeletal elements in (A) highlighting thick filaments in red, MTs in yellow and actin in light blue. The PM is in dark blue. Panels C-E show additional examples of thick filaments (arrows) and MTs (arrowheads).

Figure S2. **Gallery of tomographic slices of endosomes.** Panels A-D show a small selection of examples of endosomes with various sized internal vesicles.

Figure S3. **Gallery of tomographic slices of multivesicular bodies and their associated vesicles.** Panels A-C show typical spherical MVBs. Panels E and F show MVBS with more elliptical profiles. Panel G is a graph showing the sphericity of the MVBs, with lower values being more elliptical and a value of 1 being a perfect sphere, plotted against their equivalent sphere diameter. The line is a linear regression showing of the data and there was no significant difference between diameter and sphericity. Panels H and I show examples of MVBs with electron dense blocks of material (arrows) along with typical spherical vesicles, present in variable amounts.

Figure S4. **Gallery of tomographic slices of multilamellar vesicles and their associated internal membrane and vesicle material.** Panels A-F show the large heterogeneity in the density and variability of the material found inside the MLVs. These likely represent degradative organelles that have engulfed of fused with other membrane-bound organelles.

Figure S5. **Gallery of tomographic slices of phagophores.** Panels A-D show a gallery of phagophores in different states of maturation starting with immature (A) to fully mature where the tips have fused to form an enclosed double membrane structure (D).

Figure S6. **Galley of tomographic slices of endoplasmic reticulum and associated internal vesicles.** Panels A-F show the variability in limiting ER membrane that contain internal vesicles. Panels G and H show ER that displays a bead-on-a-string appearance.

Figure S7. **Gallery of tomographic slices of mitochondria.** The examples in panels A and B are free of electron dense precipitate, while panels C-F show evidence of precipitate.

Figure S8. **Gallery of tomographic slices showing mitochondrial remodeling.** Panels A-F show a collection of tomographic slices where mitochondria are undergoing remodeling in their transitions between the varicosity and axonal compartments. Note that several exhibit electron dense precipitate as noted in the Fig. 6B and 6C.

Movie S1 **– Movie showing the complete tomographic reconstruction corresponding to the 2D slice shown in Figure 1C**.

Movie S2 **– Complete tomographic reconstruction corresponding to the 2D slice shown in Figure 1F**.

Movie S3 **– Animation showing the 3D structure and spatial relationship of a phagophore and surrounding ER.** The outer membrane of the phagophore is in purple, the inner membrane of the phagophore is in red, cytoplasmic vesicles captured by the phagophore are in light pink, and the ER is in in dark pink.

Movie S4 **– Complete tomographic reconstruction corresponding to the 2D slice shown in Figure 6D**.

