## Supplementary Figures for "Spatial confinement shapes organelle architecture and remodeling in axons"

Thick filaments - Supplementary Figure 1

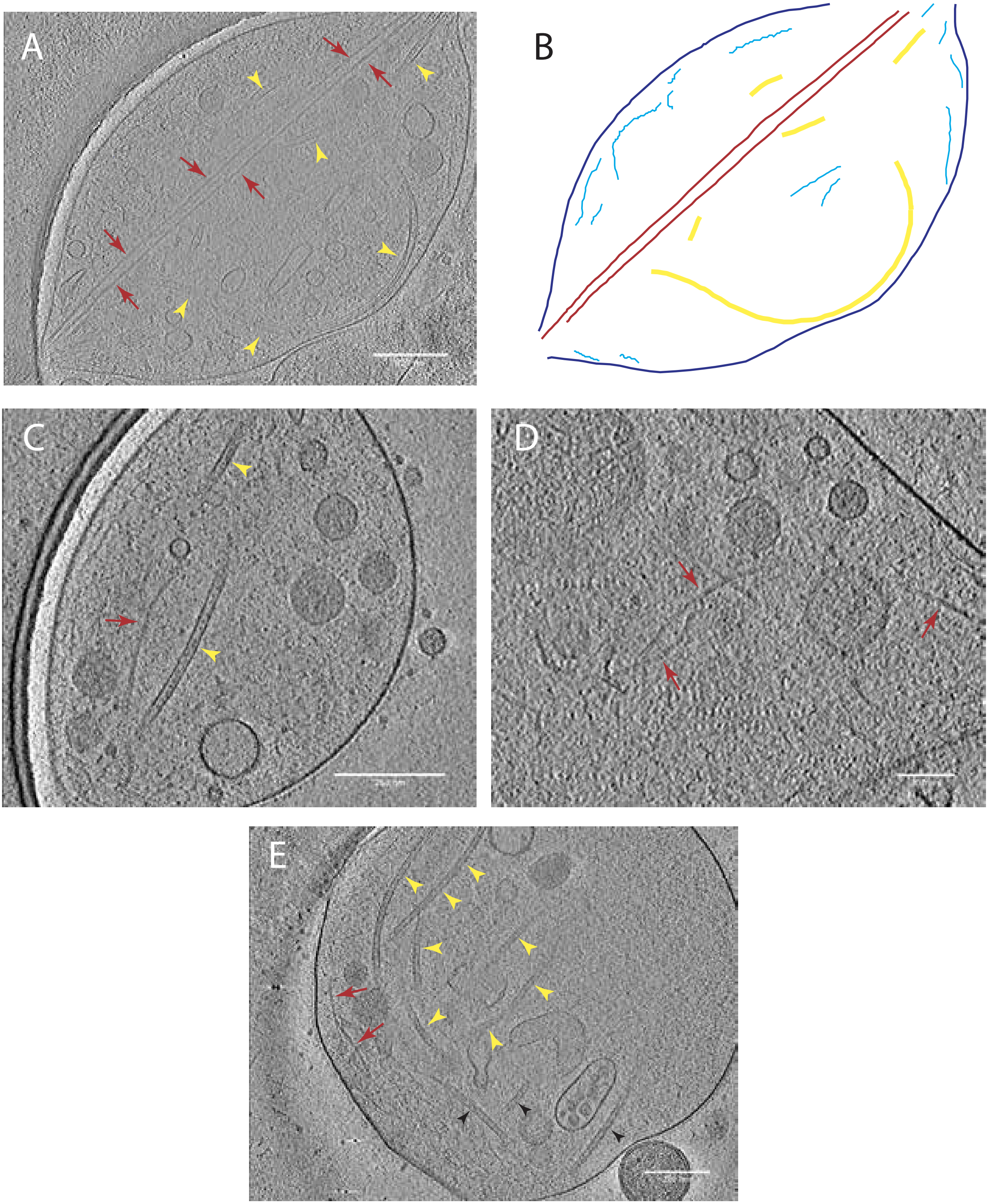

Endosomes - SI Figure 2

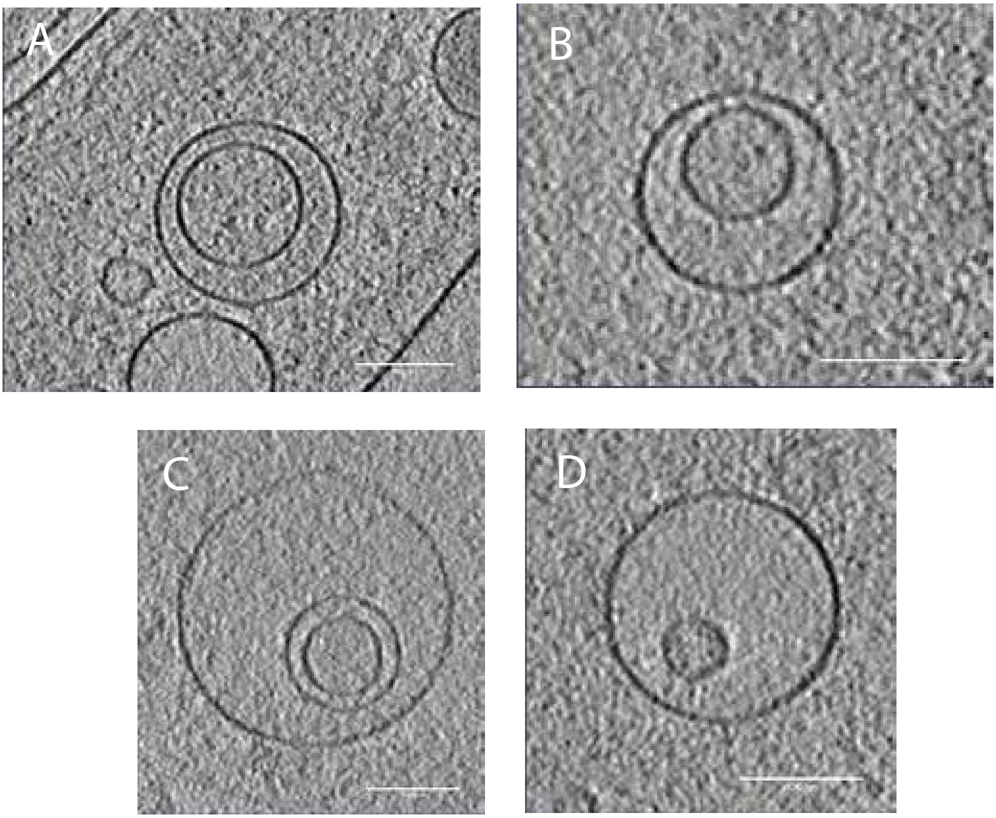

### MVBs - Supplementary Figure 3

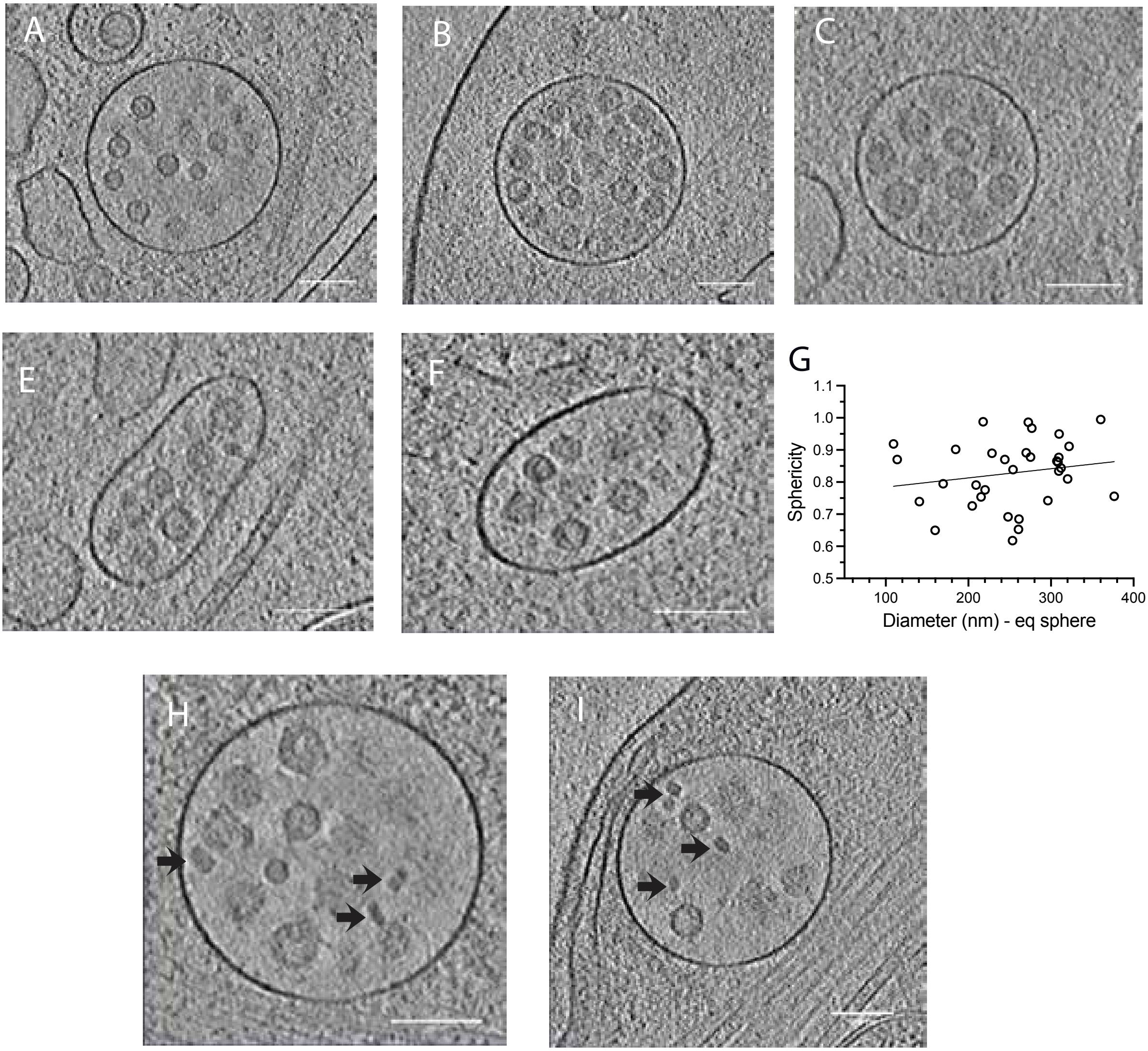

MLVs - Supplementary Figure 4

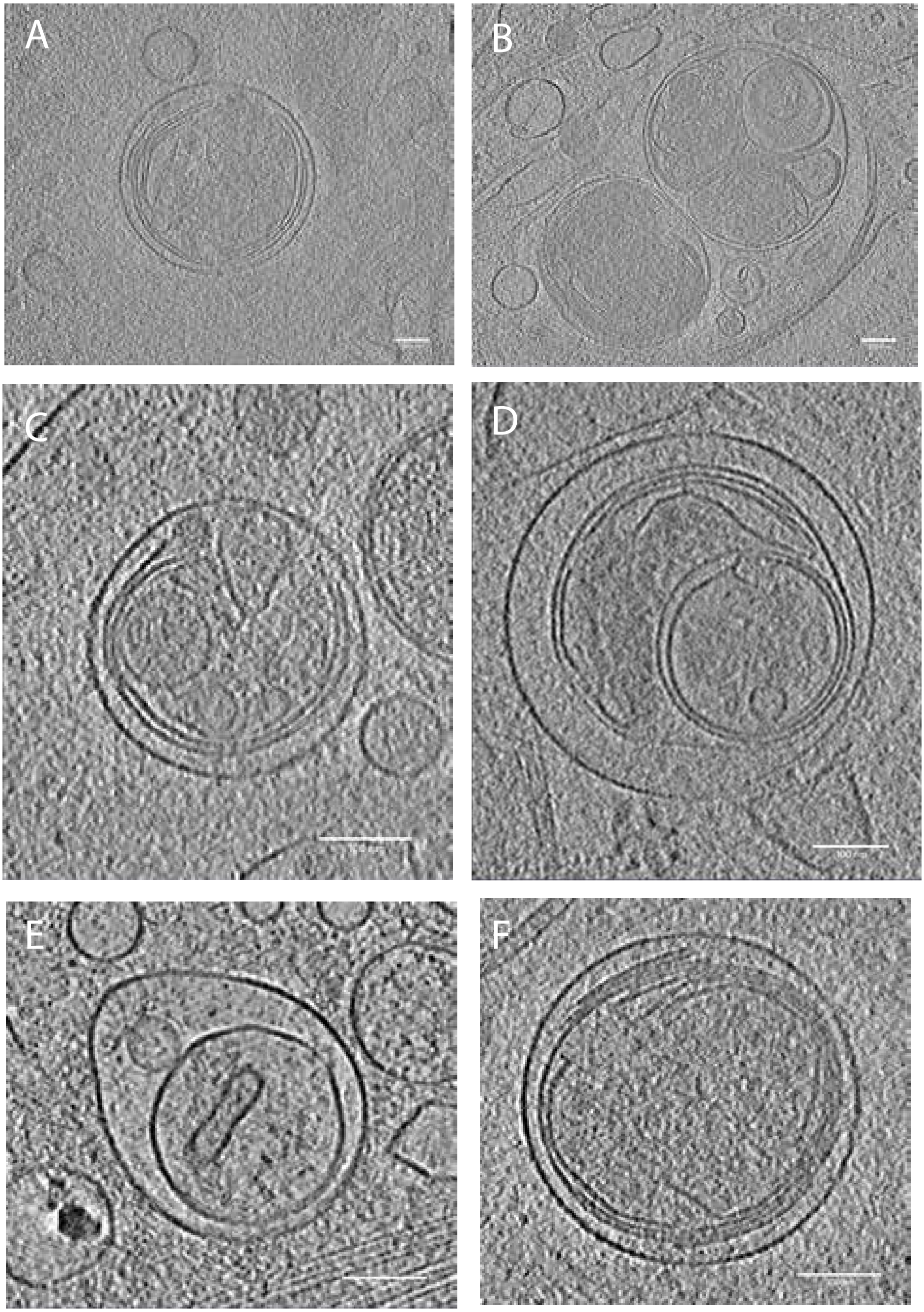

Phagophores - Supplementary Figure 5

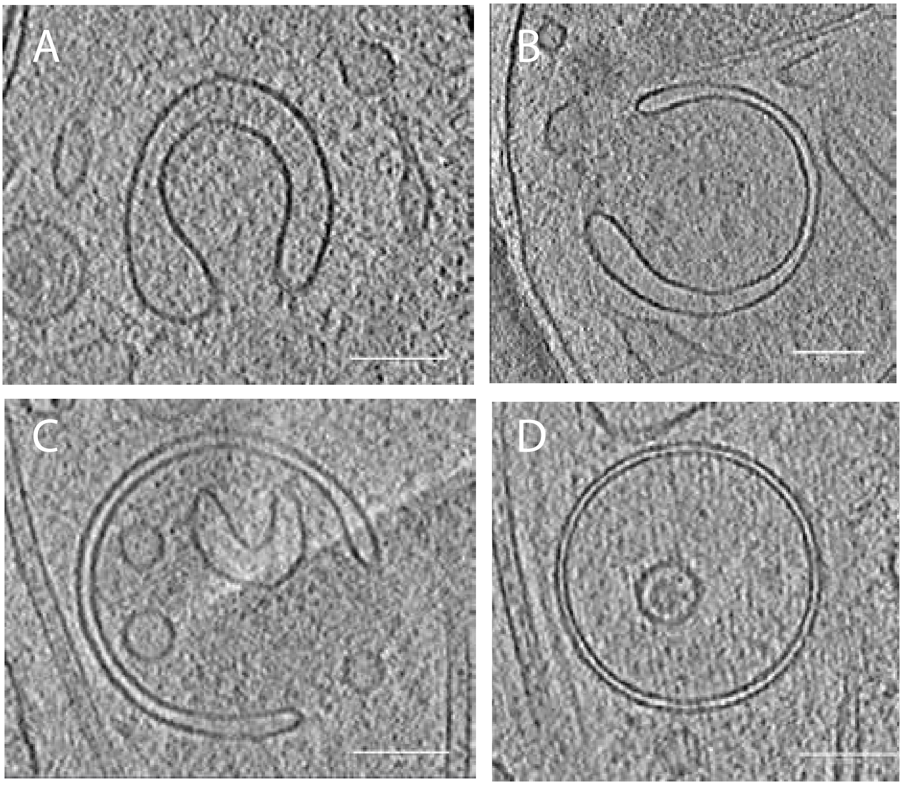

ER-Ves - Supplementary Figure 6

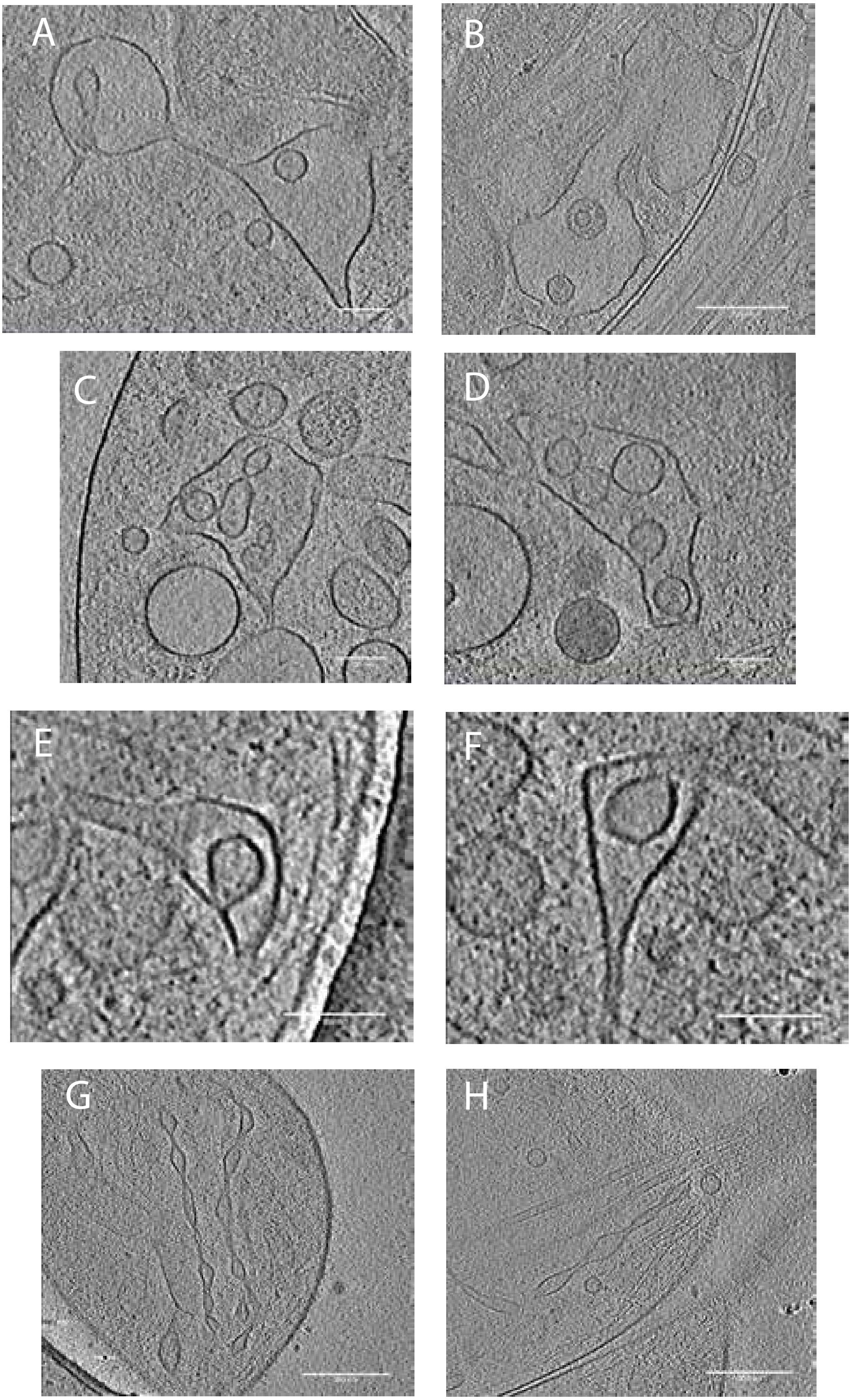

Mitochondria - Supplementary Figure 7

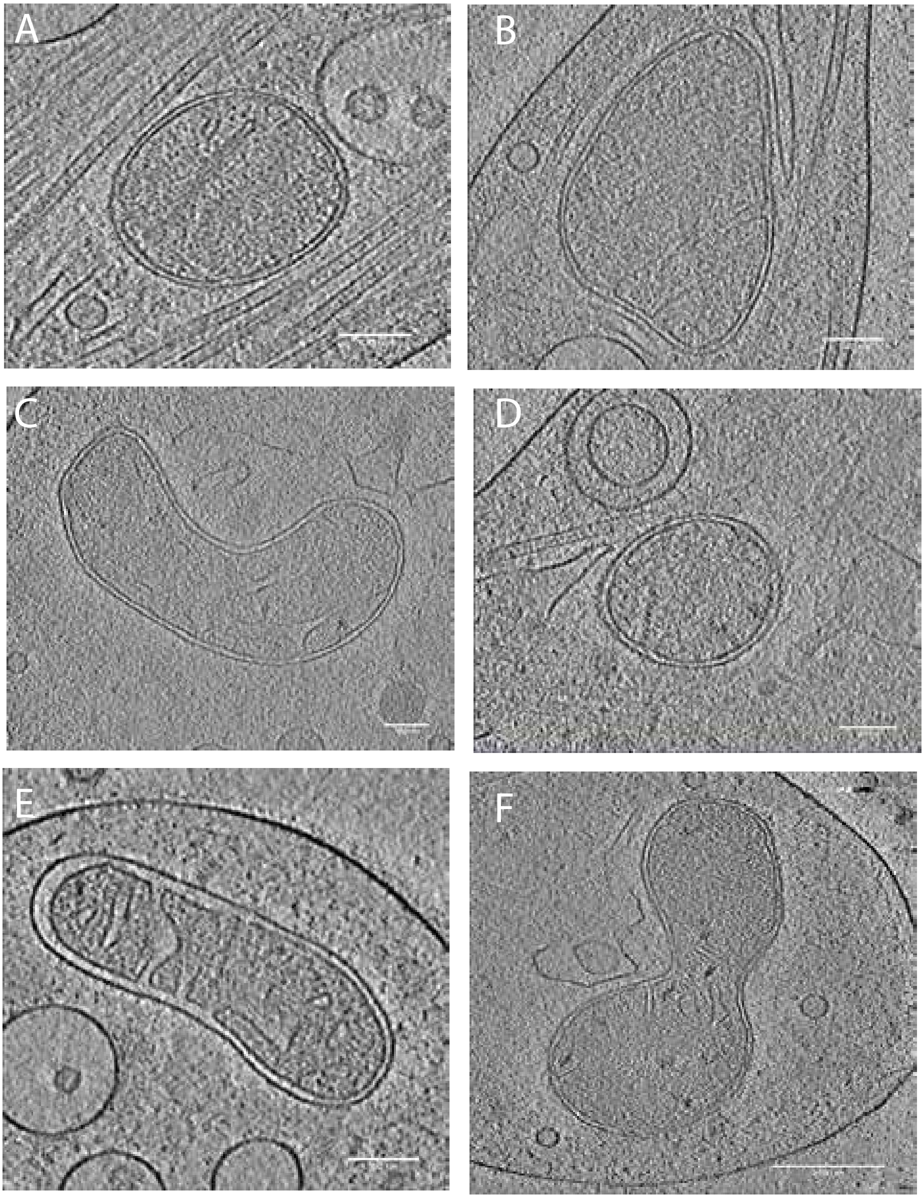

Mitochondrial Remodeling - Supplementary Figure 8

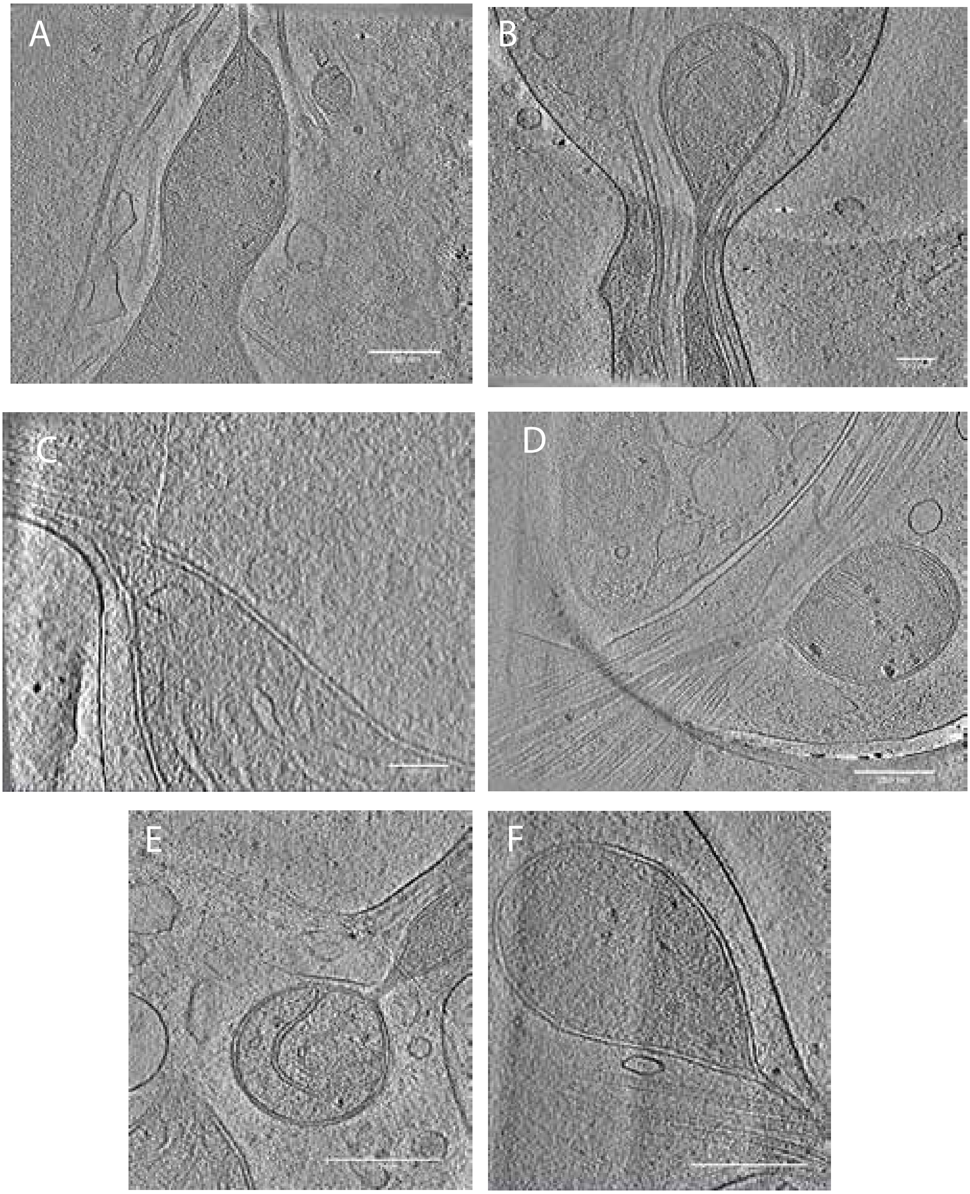
